# Regulation of Decellularized Tissue Remodeling via Scaffold-Mediated Lentiviral Delivery in Anatomically-Shaped Osteochondral Constructs

**DOI:** 10.1101/261792

**Authors:** Christopher R. Rowland, Katherine A. Glass, Adarsh R. Ettyreddy, Catherine C. Gloss, Jared Matthews, Nguyen P.T. Huynh, Farshid Guilak

**Affiliations:** Washington University in Saint Louis, Saint Louis, MO 63110; Shriners Hospitals for Children – St. Louis, St. Louis MO 63110; Duke University, Durham, NC 27710

**Keywords:** osteoarthritis, cartilage repair, regenerative medicine, immunoengineering, gene activated matrix, gene therapy

## Abstract

Cartilage-derived matrix (CDM) has emerged as a promising scaffold material for tissue engineering of cartilage and bone due to its native chondroinductive capacity and its ability to support endochondral ossification. Because it consists of native tissue, CDM can undergo cellular remodeling, which can promote integration with host tissue and enables it to be degraded and replaced by neotissue over time. However, enzymatic degradation of decellularized tissues can occur unpredictably and may not allow sufficient time for mechanically competent tissue to form, especially in the harsh inflammatory environment of a diseased joint. The goal of the current study was to engineer cartilage and bone constructs with the ability to inhibit aberrant inflammatory processes caused by the cytokine interleukin-1 (IL-1), through scaffold-mediated delivery of lentiviral particles containing a doxycycline-inducible IL-1 receptor antagonist (IL-1Ra) transgene on anatomically-shaped CDM constructs. Additionally, scaffold-mediated lentiviral gene delivery was used to facilitate spatial organization of simultaneous chondrogenic and osteogenic differentiation via site-specific transduction of a single mesenchymal stem cell (MSC) population to overexpress either chondrogenic, transforming growth factor-beta 3 (TGF-β3), or osteogenic, bone morphogenetic protein-2 (BMP-2), transgenes. Controlled induction of IL-1Ra expression protected CDM hemispheres from inflammation-mediated degradation, and supported robust bone and cartilage tissue formation even in the presence of IL-1. In the absence of inflammatory stimuli, controlled cellular remodeling was exploited as a mechanism for fusing concentric CDM hemispheres overexpressing BMP-2 and TGF-β3 into a single bi-layered osteochondral construct. Our findings demonstrate that site-specific delivery of inducible and tunable transgenes confers spatial and temporal control over both CDM scaffold remodeling and neotissue composition. Furthermore, these constructs provide a microphysiological, *in vitro*, joint, organoid model with site-specific, tunable, and inducible protein delivery systems for examining the spatiotemporal response to pro-anabolic and/or inflammatory signaling across the osteochondral interface.

## Introduction

Articular cartilage provides a smooth, low-friction bearing surface for supporting joint loading and movement. However, adult cartilage is avascular and possesses a small resident cell population of chondrocytes. These characteristics limit the capacity of cartilage to self-repair, and untreated defects ultimately progress to the full joint disease of osteoarthritis. While tissue-engineering approaches have attempted to repair defects with functional cartilage replacements, the poor intrinsic ability of cartilage to remodel prevents integration of neocartilage with the host tissue [1]. Strategies have sought to improve tissue integration by anchoring tissue-engineered constructs to the underlying subchondral bone, which possesses a strong propensity to remodel and integrate with the surrounding host tissue [2]. Fabricating osteochondral constructs with both cartilaginous and osseous phases has improved *in vivo* outcomes in rat [3] and rabbit [4, 5] animal models compared to constructs possessing either phase alone. Furthermore, subchondral bone formation has been shown to be critical for proper cartilage healing [3]. In this regard, the spatial organizing of cartilage and bone production simultaneously within osteochondral constructs has been pursued by varying site-specific scaffold composition [4–11], cell types [9, 10, 12, 13], growth factor presentation [5, 11, 13, 14], and delivery of DNA encoding for tissue-specific transcription factors [3] or growth factors [4, 8, 15].

One of the major challenges in developing integrated osteochondral constructs is the potential interference or crosstalk of signaling between the cartilaginous and osseous phases. For example, chondrogenic and osteogenic growth factors exert antagonistic effects on one another [16], and co-delivery of genes for transforming growth factor-beta 3 (TGF-β3) and bone morphogenetic protein 2 (BMP-2) reduced calcium deposition compared to either gene individually [15]. Additionally, spatially guided, dual growth factor delivery for osteochondral repair did not improve *in vivo* cartilage repair, but yielded a synergistic effect on subchondral bone formation [14]. Also, despite culturing chondrocytes and bone marrow-derived mesenchymal stem cells (MSCs) in physically separate layers of osteochondral constructs, the presence of chondrocytes inhibited bone formation by MSCs [12, 17]. This interference from competing morphogenetic signals on a single cell source such as MSCs, which can undergo both chondrogenesis and osteogenesis [4, 5, 8, 15], has encouraged the implementation of two-part scaffolds [4, 9, 10, 18–20], which are cultured in respective chondrogenic and osteogenic conditions and then later combined into a single osteochondral construct. A limitation of current two-part scaffold approaches is their reliance on external bonding techniques such as sutures [19] or fibrin glue [4, 9, 18, 20] to promote adhesion of the cartilage and bone layers. Ideally, the osteochondral interface would be stabilized through the deposition of newly synthesized matrix from seeded cells as opposed to the implementation of fixation materials, whose degradation rates might not match the rate of neotissue deposition [18].

The degradation properties not only of fixation materials but also of the scaffold itself are crucial in defining the success of osteochondral constructs upon implantation [6]. While synthetic scaffolds can offer highly tunable degradation properties, their degradation is often decoupled from cellular remodeling processes and instead is mediated by hydrolysis [3, 21, 22]. Overly rapid degradation can lead to premature loss of scaffold properties before mechanically competent tissue has formed, resulting in implant failure [23]. Conversely, the persistence of synthetic polymers beyond the remodeling process can induce a foreign body response [24] and fibrotic tissue deposition [3, 14, 25, 26], which inhibit subchondral bone remodeling [3, 14, 26] and cause osseous wall resorption and defect widening [6]. Additionally, non-degradable hydrogels sequester newly synthesized matrix to the pericellular region [27, 28], and implants delivering TGF-β3 and BMP-2 only observed tissue elaboration in degradable scaffolds [27], which highlights the importance of scaffold degradation in osteochondral repair. While the incorporation of enzymatically-cleavable degradation sites within synthetic polymers have associated scaffold degradation with the remodeling process [28, 29], there is still concern over the degradation products of synthetic scaffolds, which stimulate macrophages to secrete inflammatory cytokines, catabolic enzymes, and cytotoxic factors [22]. In contrast, scaffolds derived from the extracellular matrix of native tissues retain the capacity to be remodeled by cells, and their degradation products promote a constructive remodeling M2 phenotype in macrophages [30, 31], which has been correlated with more favorable outcomes in preclinical animal models [32].

Interestingly, in the absence of exogenous growth factors, degradation products of the cartilage extracellular matrix possess the capacity to stimulate chondrogenic differentiation of MSCs in a dose-dependent manner [33]. Furthermore, even in the presence of TGF-β inhibitors, cartilage-derived matrix (CDM) activates chondrogenic genes and exhibits a potentiated response with TGF-β3 supplementation, suggesting that the mechanisms by which CDM induces chondrogenic differentiation are independent of and synergistic with exogenous growth factors [34]. Not only has CDM been used extensively as a biomaterial for cartilage regeneration [35], but it also has been incorporated into biphasic constructs for *in vivo* osteochondral repair [36–38]. Within biphasic constructs, CDM was only used in the cartilage portion, while the osseous phase was fabricated with materials that mimicked the composition of bone to promote osteogenesis through the intramembranous ossification pathway [36–38]. However, intramembranous ossification generates excessive mineralization that inhibits vascularization of the subchondral bone [12] producing a necrotic core [39], which could be problematic for larger implants. In contrast, endochondral ossification recapitulates the developmental process of bone formation, and utilizes a cartilaginous substrate for generating mature mineralized tissue with a vascular network [40]. These benefits highlight the spatial regulation of endochondral ossification as a promising strategy for fabricating osteochondral constructs [9, 12, 15]. CDM has been used as a template for endochondral bone formation [41]. Therefore, the ability of CDM to support both cartilage [35] and bone [41] regeneration empowers CDM to serve as a single, compositionally homogenous substrate for engineering osteochondral constructs. Additionally, CDM can be remodeled *in vivo* [42]. However, the *in vivo* degradation profile of CDM is uneven, random, and irregular over time [43], which contributes to the failure of CDM constructs in long-term, high load bearing, osteochondral repair [44]. Attempts to delay the degradation of CDM have included the use of chemical crosslinking agents, which confer resistance to enzymatic degradation [45, 46]; however, chemically-crosslinked, tissue-derived scaffolds can evoke chronic foreign body response, fibrotic encapsulation, and ultimately poor outcomes *in vivo* [32]. Moreover, chemical crosslinking treatments negatively impact the ability of CDM to participate in cell-matrix interactions, and diminish the chondroinductive capacity of CDM [47]. Therefore, there is a need to be able to tailor scaffold degradation without modifying the intrinsic properties of CDM via chemical crosslinking.

The overarching goal of this study was to spatially and temporally control both scaffold degradation properties as well as simultaneous cartilage and bone formation within anatomically-shaped CDM scaffolds. As opposed to implementing chemical crosslinking techniques, the current study utilized dehydrothermal treatment, which is a physical crosslinking method that increases the compressive modulus of CDM scaffolds [48] and prevents cell mediated contraction [49], while still preserving cell-matrix interactions [47] and enzymatic remodeling [50]. In order to regulate scaffold degradation, we examined the hypothesis that IL-1Ra production can be used to control the degradation of MSC-seeded CDM scaffolds treated with IL-1 at levels found in the OA joint [51]. Specifically, interleukin-1 (IL-1) receptor antagonist (IL-1Ra) has been shown to decrease inflammatory mediators and catabolic proteases in an *in vivo* model of primary OA [52] and to influence macrophage polarization towards a constructive remodeling M2 phenotype *in vitro* [53]. Furthermore, viral gene delivery of IL-1Ra has been shown to slow the rate of cartilage loss *in vivo* [54] and prevented IL-1-mediated atrophy of tissue-engineered cartilage [55]. Therefore, controlled IL-1Ra production may inhibit cell-based catabolic processes and thus reduce the CDM degradation rate. To spatially dictate protein secretion, scaffold-mediated gene delivery was utilized because it overcomes the limitations of protein immobilization such as a burst-release profile, short half-lives of active forms, and restrictions on the overall quantity of protein available [56–58]. While non-viral [57] and adeno-associated viral [59–61] gene delivery may face obstacles of low transfection efficiency and transient gene expression [56], scaffold-mediated lentiviral gene delivery yields high transduction efficiency and long-term, stable expression [55, 62, 63]. Additionally, doxycycline (dox)-inducible promoters provide control over both the magnitude and duration of expression [55]. Here we demonstrate that immobilization of lentivirus containing either TGF-β3 or BMP-2 transgenes in spatially distinct regions of anatomically-shaped CDM scaffolds [49] allows for the simultaneous formation of cartilage and bone constructs in a site-specific manner, which in addition to lentiviral gene delivery of IL-1Ra, offers spatial and temporal control over both CDM scaffold remodeling and tissue-specific matrix deposition. In addition to direct application to tissue engineering, these constructs provide an *in vitro* model of osteochondral organoids that possess site-specific tunable and inducible protein delivery systems for microphysiological testing.

## Methods

### CDM Hemisphere Fabrication

Porcine articular cartilage was harvested, decellularized, and pulverized into a fine powder as described previously [49]. CDM powder was suspended in water via homogenization at a concentration of 11% w/w and pipetted into the following delrin molds with silicone lids: 1) hemispherical shells with an outer radius of 4.76 mm and inner radius of 3.175 mm, and 2) hemisphere cores with a radius of 3.175 mm. To make the hemispherical shells, the silicone lid for the 4.76 mm diameter hemispheres had hemisphere protrusions of 3.175 mm radius. CDM hemispheres were then frozen and lyophilized as described previously [49]. All CDM hemispheres were minimally crosslinked and sterilized via dehydrothermal treatment (120°C for 24hrs).

### Lentivirus Production and Immobilization on CDM Hemispheres

Doxycycline-inducible lentiviral vectors were generated by cloning eGFP, TGF-β3, BMP-2, or IL-1Ra coding sequences into the modified TMPrtTA vector (provided by the Danos Lab) as described previously [55, 62] (see Supplemental Fig. 1 for vector map). Lentivirus (LV) was concentrated ~75-fold using Amicon Ultra 100 kDa MWCO filters (Millipore, Cork, Ireland) and frozen at −80°C until use. To immobilize LV, CDM hemispheres were incubated overnight with 0.002% poly-L-lysine (PLL, Sigma-Aldrich) in PBS. Hemispheres were washed twice in PBS prior to seeding 150 µL of concentrated LV, which was allowed to attach for 1.5 hrs at 37°C. To remove non-adherent LV, hemispheres were rinsed twice with PBS and either immediately seeded with cells (Fresh Virus) or frozen and lyophilized (Freeze Dried) prior to cell seeding. While the LV freeze dried onto CDM hemispheres was capable of transducing cells, the transduction efficiency and resultant protein secretion were significantly lower compared to freshly immobilized LV (Supplemental Fig. 2). For the current study, all CDM hemispheres were seeded with cells immediately following LV immobilization (fresh virus).

**Figure 1:**
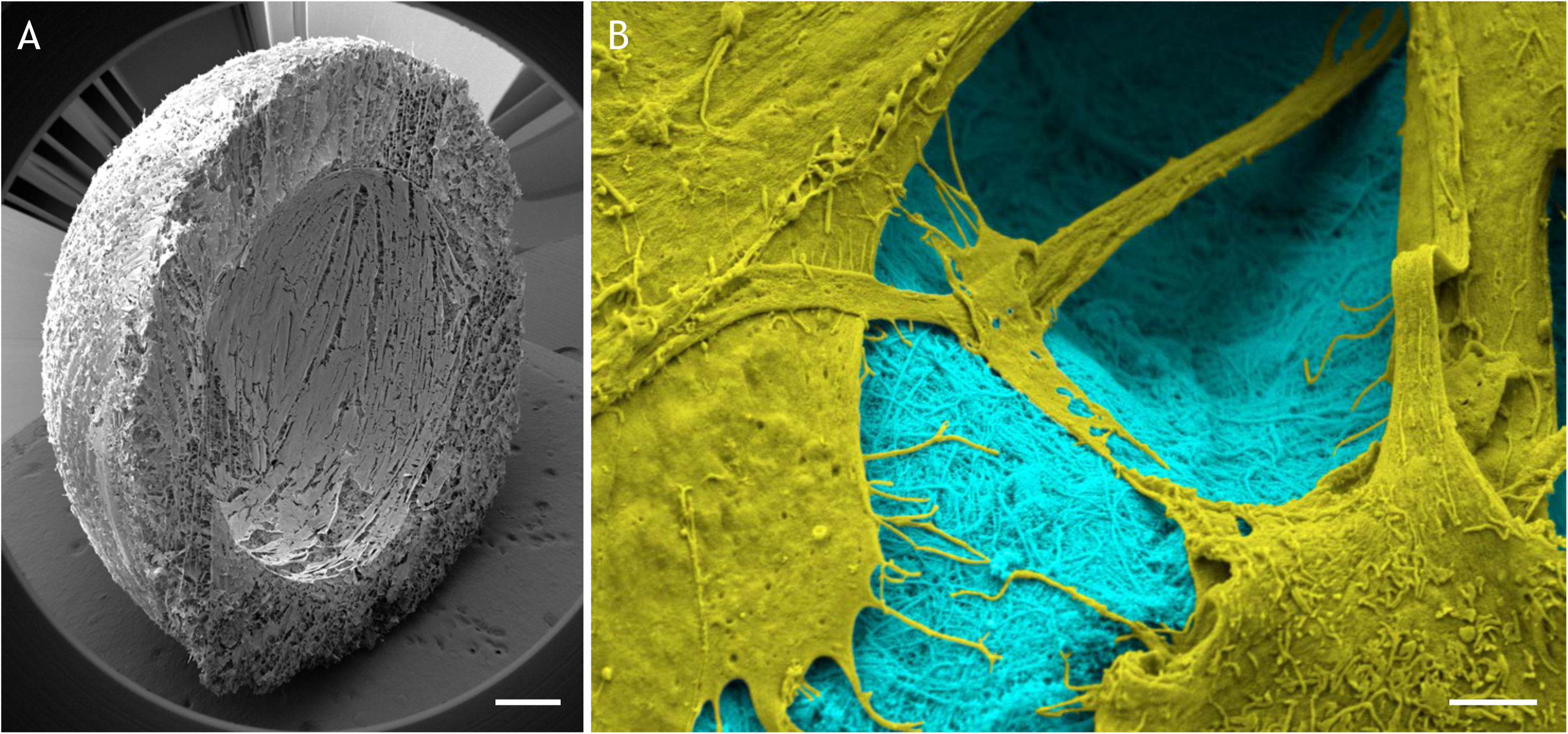
Low (A) and high (B) magnification scanning electron microscope images of a CDM hemisphere (blue) with immobilized eGFP lentivirus 1 hr after seeding MSCs (yellow). (A) Scale: 1mm. (B) Scale: 2μm.

### Cell Culture

Bone marrow aspirate was obtained from three de-identified adult transplant donors at Duke University Medical Center, in an IRB-exempt use of waste tissue. Human MSCs were isolated, expanded, and pooled into a superlot as described previously [49]. Passage 4 MSCs were trypsinized, counted, and suspended at a concentration of 6.67e6 cells/mL. A cell suspension (150 μL) was seeded onto PLL-coated CDM hemispheres without LV (non-transduced) or immobilized with eGFP, TGF-β3, BMP-2, or IL-1Ra LV. Cells were allowed to attach for 1 hr at 37°C prior to adding 2 mL of expansion medium as defined previously [49]. After 6 days of cell expansion on CDM hemispheres, constructs were switched to one of the following media conditions: basal chondrogenic, basal osteogenic, chondrogenic with 10 ng/mL rhTGF-β3 (R&D Systems, Minneapolis, MN), osteogenic with 100 ng/mL rhBMP-2 (R&D Systems), chondrogenic with 10 ng/mL rhTGF-β3 and 0.1 ng/mL rhIL-1α (R&D Systems), or osteogenic with 100 ng/mL rhBMP-2 and 0.1 ng/mL rhIL-1α. Basal chondrogenic medium consisted of DMEM-HG (Gibco), 1% Penicillin-Streptomycin (Gibco), 1% ITS^+^ (Corning), 100 nM dexamethasone (Sigma-Aldrich), 50 µg/ml L-ascorbic acid (Sigma-Aldrich), and 40 µg/ml L-proline (Sigma-Aldrich). Basal osteogenic medium consisted of DMEM-HG, 10% lot-selected FBS (HyClone, Thermo Scientific, Waltham, MA), 1% Penicillin-Streptomycin, 50 µg/mL L-ascorbic acid, 10 nM dexamethasone, and 10 mM β-glycerol phosphate. Each of the media conditions was supplemented with 1 μg/mL doxycycline (dox) to induce transgene expression, and switching to the respective media conditions corresponded with day 0 of differentiation. rhIL-1α (0.1 ng/mL) was added at day 2 of respective culture conditions. Constructs were cultured for 28 days with half-media changes every 48 hrs. Media were collected at days 0, 2, 6, 10, 16, 22, and 28 and stored at −20°C for analysis. Constructs were harvested at 1 hr after cell seeding, day 0 (after 6 days of expansion), and day 28.

### Osteochondral Construct Fabrication

As described above, TGF-β3 LV was immobilized onto outer hemispherical shells, while BMP-2 LV was immobilized onto inner hemisphere cores (n=8). For control samples, eGFP LV was immobilized on both hemispherical shells and hemisphere cores (n=4). Hemispherical shells were seeded with 150 μL of a 6.67e6 cells/mL suspension of passage 4 MSCs, while hemisphere cores were seeded with 75 μL of the same cell suspension. Following a 6-day expansion period, hemispherical shells were cultured in basal chondrogenic medium and hemisphere cores were cultured in basal osteogenic media for 22 days. After respective chondrogenic and osteogenic culture, hemisphere cores were press-fit into hemispherical shells (**Supplemental Video**) and cultured together in basal chondrogenic media for an additional 28 days. For the entire differentiation period, samples were treated with 1 µg/mL dox and half of the media was replaced every 48 hrs. Media was collected at days 0, 6, 16, 22, 28, 38, and 50. Constructs were harvested at day 50.

### Scanning Electron Microscopy

One hour after cell seeding, CDM constructs were fixed, dehydrated, critical point dried, and sputter-coated as described previously [49]. Samples were imaged at 3 kV, 90 pA, 11.16 nm pixel size using a Crossbeam 540 Focused Ion Beam Scanning Electron Microscope (Zeiss). Images were false-colored in Adobe Photo Shop.

### Transduction Efficiency

Twenty days after scaffold-mediated transduction, non-transduced (NT) and eGFP-transduced constructs (n=3/group) were cut in half and cross-sections were imaged using a confocal microscope (LSM 510, Zeiss). Gain was set such that there was no background in the NT constructs and the same gain was applied for all samples. After imaging, constructs were digested in 1320 PKU/mL Pronase (Calbiochem, San Diego, CA) in DMEM-HG with 5% FBS for 1 hr at 37°C followed by subsequent digestion in 0.002 g/mL type II collagenase (Worthington, Lakewood, NJ) and 10 µg/mL DNase I (Sigma, St. Louis, MO) for 2.5 hrs at 37°C with gentle shaking. Isolated MSCs were filtered through 70 µm cell strainers, washed twice in PBS, and fixed in 1% PFA in PBS. Flow cytometry was performed to obtain the percentage of eGFP+ cells using the Accuri flow cytometer (BD Biosciences, Franklin Lakes, NJ) with 10,000 events counted per sample.

### Media Analyses

Enzyme-linked immunosorbent assays (ELISAs) for human TGF-β3, BMP-2, and IL-1Ra were performed on media from various time points according to the manufacturer’s instructions (n=4–8/group) (R&D Systems). Samples were run in duplicate and optical density was measured at 450 nm with a correction at 540 nm. Latent TGF-β3 was activated by a 10-min incubation with HCl. The exogenous rhTGF-β3 added to the media was also measured as a control. Total specific MMP activity was quantified as described previously [64].

### Biochemical Quantification

Constructs (n=6/group) were lyophilized and then digested in papain buffer [125 μg/mL papain (Sigma), 100 mM phosphate buffer, 10 mM cysteine, and 10mM EDTA, pH 6.3] for 16 hrs at 65°C. As described previously [49], orthohydroxyproline assay [65], dimethylmethylene blue (DMMB) assay [66], and Quant-iT™ PicoGreen^®^ dsDNA Assay Kit (Invitrogen) were used to quantify total collagen, glycosaminoglycan (GAG), and double-stranded DNA content, respectively. Calcium content was measured in the decal solution from histology samples using a colorimetric assay (n=2/group) (BioVision, Milpitas, CA).

### Histology and Immunohistochemistry

Prior to fixation, all constructs in osteogenic media and three chondrogenic negative controls were decalcified in 2 mL of 5% formic acid in deionized water for 30 min at room temperature. Decal solution was collected and frozen at −20°C for calcium quantification. Constructs (n=2/group) were fixed, dehydrated, cleared, paraffin embedded, and sectioned (8 µm thickness) as described previously [49]. Immunohistochemistry (IHC) was performed with monoclonal antibodies against type I collagen (ab90395; Abcam, Cambridge, MA), type II collagen (II-II6B3; Developmental Studies Hybridoma Bank, University of Iowa), or type X collagen (C7974; Sigma-Aldrich) using aminoethyl carbazole (Invitrogen) as the chromogen as described previously [49]. Xylene-cleared sections were also histologically stained using 0.1% aqueous Safranin-O, 0.02% fast-green, and hematoxylin. Human osteochondral tissue was used as a positive control. Negative controls for IHC did not include primary antibody. Sections were imaged on a CKX41 scope with a DP26 camera (Olympus).

### Micro-Computed Tomography Analysis

CDM constructs (n=6/group) were scanned using micro-computed tomography (micro-CT) (SkyScan 1176, Bruker, Billerica, MA) at 40 kV, 555 μA, and 17.51 μm isotropic spatial resolution. Micro-CT datasets were reconstructed with NRecon software (Bruker) using a dynamic range of 0.09, ring artifact correction of 11, and beam hardening correction of 20%. Mineralization was quantified via calibration against hydroxyapatite phantoms using CT-Analyzer software (Bruker). Mineral distribution ratio was calculated by dividing the mineral volume in the bottom half of CDM constructs by the mineral volume in the top half. All hemispherical constructs were cultured in the same orientation with their convex portion resting on the bottom of tissue culture dishes. The bottom and top halves represent equal volumes of the hemispherical scaffold with the “bottom” referring to the half of the scaffold that was resting on the tissue culture surface while the “top” half was furthest away from the tissue culture surface.

### Statistical Analysis

Statistical analyses were performed using JMP Pro 12 (SAS Institute Inc., Cary, NC). Differences between two groups were determined using a student’s t-test (p<0.05). Differences between groups with media condition and vector condition as factors were determined using two-way ANOVA with Tukey’s post-hoc analysis (α=0.05).

## Results

Scanning electron microscopy (SEM) illustrated the concave and convex architecture as well as the porous nature of the CDM hemispheres (Fig. 1A), and revealed that MSCs readily adhered to PLL-coated CDM hemispheres with immobilized lentivirus (Fig. 1B). After 6 days of expansion on CDM hemispheres, which corresponded with day 0 of differentiation, MSCs completely infiltrated the entire thickness of CDM constructs (Supplemental Fig. 3A). Additionally, constructs maintained their hemispherical shape and no discernible matrix accumulation was seen during the expansion period (Supplemental Fig. 3B).

ELISA on collected media showed that protein secretion was inducible and stable throughout the culture period (Fig. 2). Doxycycline (dox) was not administered during the 6-day expansion culture and was supplemented beginning at day 0 of differentiation. In the absence of dox, the baseline TGF-β3 (Fig. 2A), BMP-2 (Fig. 2B), and IL-1Ra (Fig. 2C) concentrations were respectively undetectable, 0.93 ± 0.06 ng/mL, and 1.66 ± 0.09 ng/mL. The negligible protein detected at day 0 followed by the sharp increases by day 6 confirmed that TGF-β3 (Fig. 2A), BMP-2 (Fig. 2B), and IL-1Ra (Fig. 2C) secretion were indeed dox-inducible. Protein synthesis was not only inducible, but also stable – TGF-β3 (Fig. 2A), BMP-2 (Fig. 2B), and IL-1Ra (Fig. 2C) were detected in the culture media throughout the 28-day differentiation period and eGFP expression was observed in all media conditions at day 28 (Supplemental Fig. 4). Critically, there was no effect of rhIL-1 treatment on IL-1Ra synthesis in either chondrogenic or osteogenic media (Fig. 2C). While osteogenic media produced a lower IL-1Ra concentration compared to chondrogenic media, there was no interaction between media type and rhIL-1 treatment on IL-1Ra secretion (Fig. 2C).

**Figure 2:**
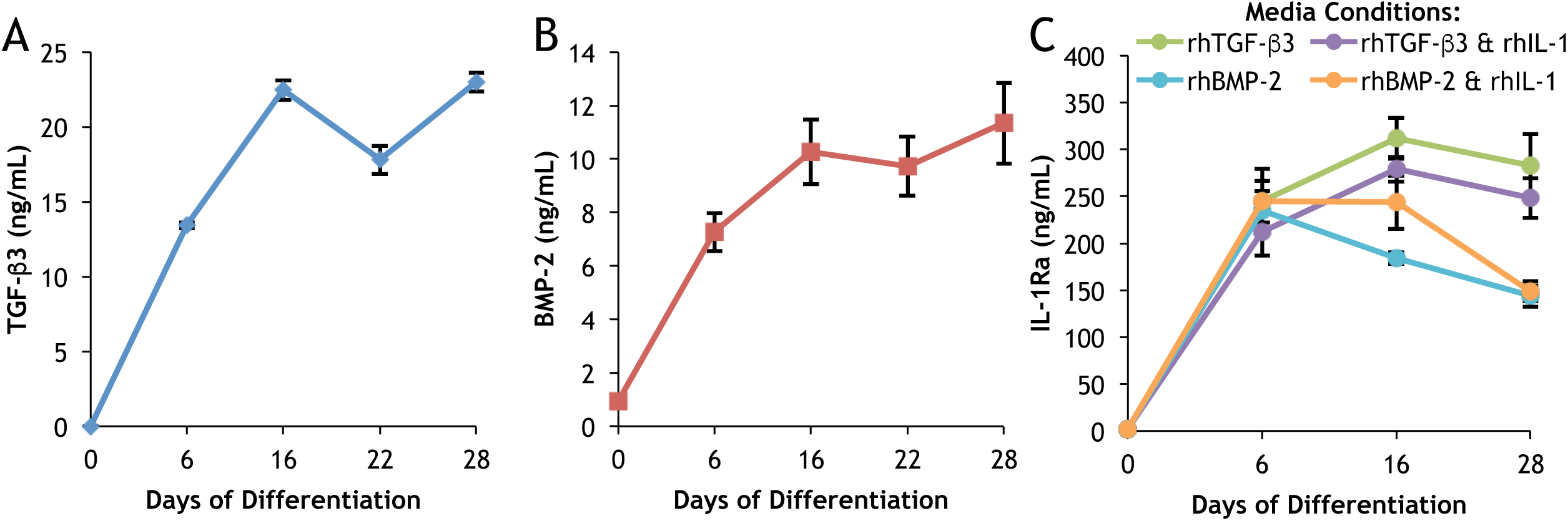
Protein secretion of CDM constructs expressing dox-inducible (A) TGF-β3, (B) BMP-2, or (C) IL-1Ra. Constructs were cultured for 28 days with 1 µg/mL doxycycline added at day 0 in (A) basal chondrogenic media (without exogenous rhTGF-β3); (B) basal osteogenic media (without exogenous rhBMP-2); (C) either chondrogenic (with rhTGF-β3) or osteogenic (with rhBMP-2) media conditions in the presence or absence of rhIL-1. Mean ± SEM (n=4). Transduction efficiency was 57 ± 5% eGFP+ MSCs (mean ± SEM, n=3).

MMP activity analysis revealed that IL-1Ra expression inhibited the catabolic effects of rhIL-1 treatment (Fig. 3). The day 2 data shows baseline MMP activity prior to rhIL-1 treatment for non-transduced (NT), eGFP-transduced, and IL-1Ra-transduced constructs (Fig. 3). At day 6, there was no effect of rhIL-1 treatment on MMP activity within NT, eGFP-transduced, or IL-1Ra-transduced constructs; however, at day 10 there was a significant effect of rhIL-1 treatment in NT constructs (Fig. 3). By day 16, rhIL-1 treatment dramatically increased MMP activity in both NT (~36-fold) and eGFP-transduced (~22-fold) constructs (Fig. 3). At day 22 and day 28 in both NT and eGFP-transduced constructs, rhIL-1 treatment continued to elevate MMP activity, which peaked at day 28 with fold increases of ~59 and ~46 for NT and eGFP-transduced constructs, respectively (Fig. 3). Interestingly, there was no upregulation of MMP activity in IL-1Ra-transduced constructs throughout the entire culture period (Fig. 3)

**Figure 3:**
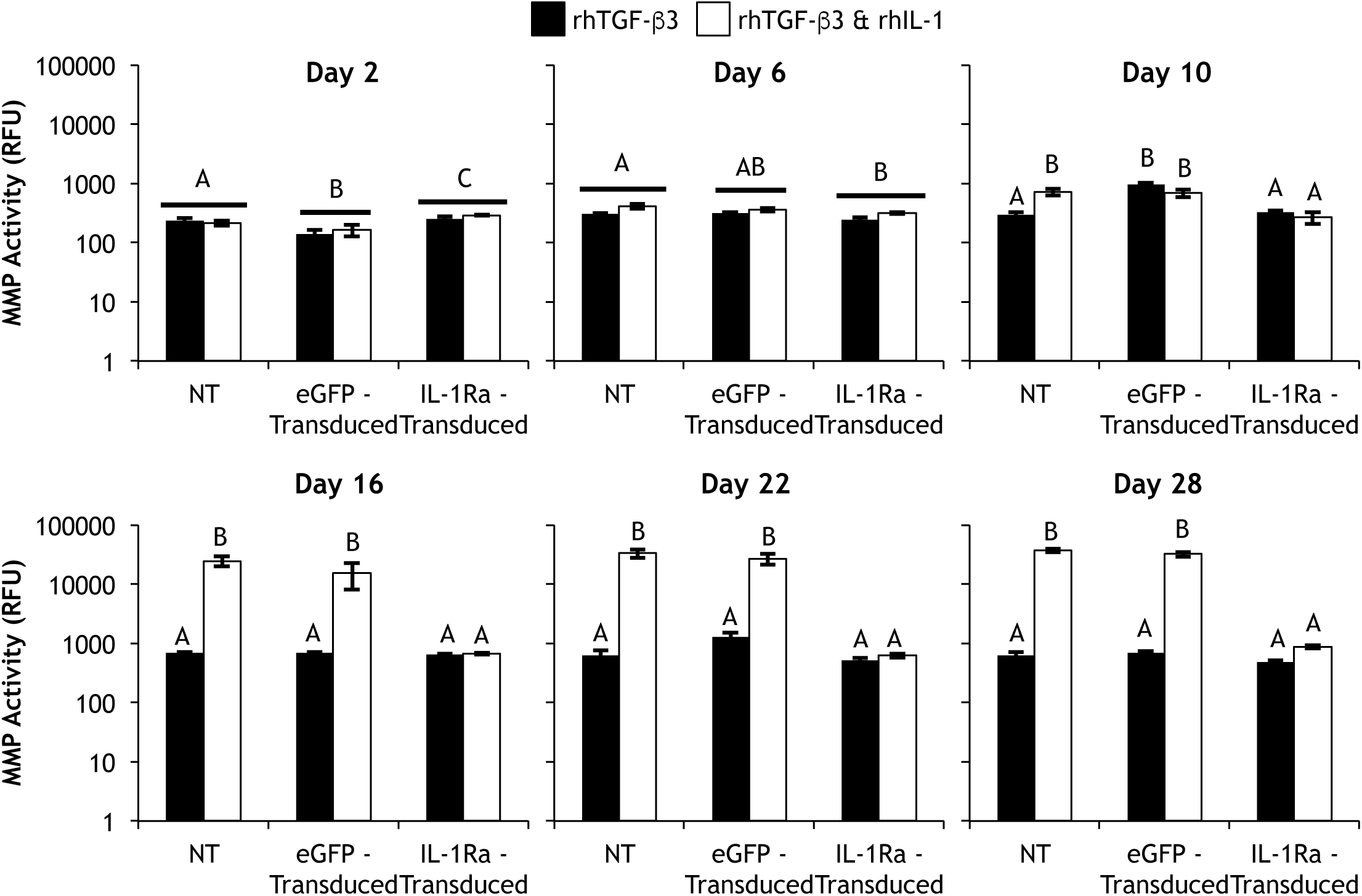
Total specific MMP activity of non-transduced (NT), eGFP-transduced, or IL-1Ra-transduced constructs cultured in chondrogenic media containing rhTGF-β3 either without or with rhIL-1 added beginning at day 2. Bars represent mean ± SEM (n=4). Groups not sharing same letter have p-values < 0.05.

Scaffold-mediated lentiviral gene delivery did not affect the baseline biochemical composition of CDM constructs; at day 0 of differentiation (after 6 days of expansion) there was no difference in the collagen, GAG, or DNA contents between NT and eGFP-transduced constructs (reported in the caption of Fig. 4). Since cartilage extracellular matrix is rich in collagen, the collagen content of cultured constructs is predominantly CDM (Fig. 4). In chondrogenic culture, neither TGF-β3 expression nor exogenous supplementation of rhTGF-β3 produced significant increases in collagen compared to NT and eGFP-transduced controls (Fig. 4A). However, in both NT and eGFP-transduced constructs, rhIL-1 treatment elicited dramatically lower collagen contents. There was no difference in the collagen content of IL-1Ra-transduced constructs in the presence or absence of rhIL-1 (Fig. 4B). Interestingly, the same trends were seen in osteogenic media (Fig. 4C & **4D**). There was no effect of BMP-2 expression or exogenous supplementation of rhBMP-2 on collagen content (Fig. 4C); and rhIL-1 treatment produced lower collagen contents in both NT and eGFP-transduced constructs but not in IL-1Ra-transduced constructs (Fig. 4D).

**Figure 4:**
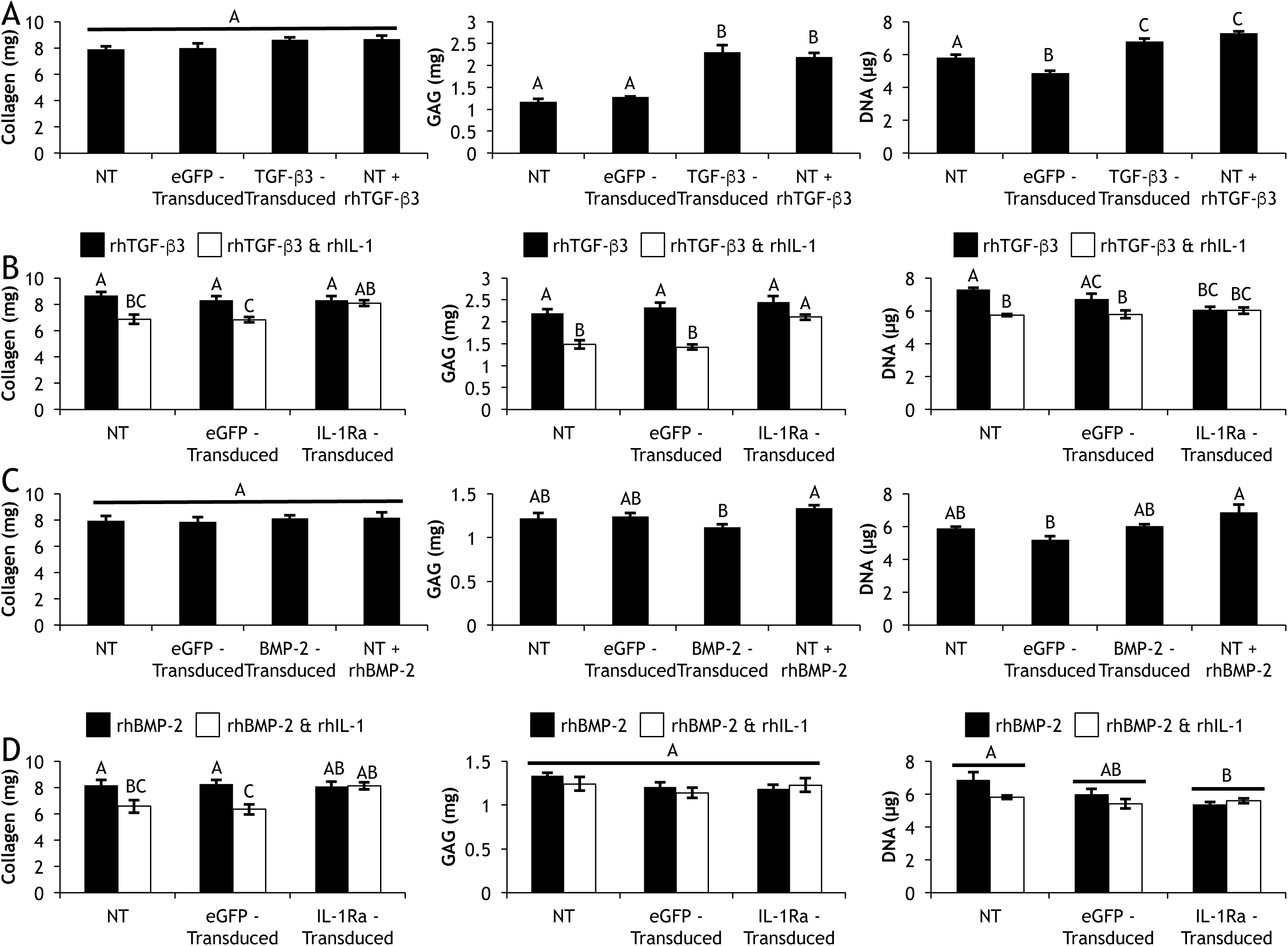
Day 28 collagen (left), GAG (middle), and DNA (right) contents of (Row A) non-transduced (NT), eGFP-transduced, or TGF-β3-transduced constructs cultured in basal chondrogenic media (without exogenous rhTGF-β3), and NT constructs fed exogenous rhTGF-β3; (Row B) NT, eGFP-transduced, and IL-1Ra-transduced constructs cultured in chondrogenic media containing rhTGF-β3 with and without rhIL-1; (Row C) NT, eGFP-transduced, or BMP-2-transduced constructs cultured in basal osteogenic media (without exogenous rhBMP-2), and NT constructs fed exogenous rhBMP-2; (Row D) NT, eGFP-transduced, and IL-1Ra-transduced constructs cultured in osteogenic media containing rhBMP-2 with and without rhIL-1. Mean ± SEM (n=6). Groups not sharing same letter have p-values < 0.05. At day 0, Collagen = 8.24mg, GAG = 2.11mg, DNA = 4.36μg.

While CDM is initially rich in proteoglycans, GAGs leached out of constructs over time and by day 28 both NT and eGFP-transduced constructs possessed less GAG than at day 0 of chondrogenic differentiation (Fig. 4A). However, either TGF-β3 expression or rhTGF-β3 supplementation resulted in significant increases in GAG content compared to NT and eGFP-transduced controls, and there was no difference in GAG content between TGF-β3 expression and rhTGF-β3 supplementation (Fig. 4A). Treatment with rhIL-1 significantly decreased GAG content in both NT and eGFP-transduced constructs; however, IL-1Ra-transduced constructs maintained GAG levels that were not different from constructs cultured in the absence of rhIL-1 (Fig. 4B). Osteogenic differentiation conditions did not lead to GAG production, and all constructs lost GAG relative to Day 0 (Fig. 4C & 4D).

Since CDM was decellularized prior to hemisphere fabrication, it did not contribute to the DNA content of cultured constructs, and the day 0 DNA content reflected the MSC population after the 6-day expansion period. In chondrogenic media, all constructs experienced cellular proliferation over the 28-day differentiation period (Fig. 4A). However, both TGF-β3-transduced constructs and constructs fed rhTGF-β3 had significantly more cellular proliferation than NT constructs, which in turn had higher proliferation than eGFP-transduced constructs (Fig. 4A). Treatment with rhIL-1 yielded lower DNA contents in both NT and eGFP-transduced constructs compared to respective constructs cultured in the absence of rhIL-1 (Fig. 4B). However, there was no difference in the DNA contents of IL-1Ra-transduced constructs cultured in the presence or absence of rhIL-1 (Fig. 4B). In osteogenic media, all constructs experienced cellular proliferation by day 28; and there was no difference in DNA content of NT, BMP-2-transduced, or constructs fed rhBMP-2 (Fig. 4C). Additionally, in osteogenic medium, there was no difference in the DNA contents of NT, eGFP-transduced, or IL-1Ra-transduced constructs cultured in the presence or absence of rhIL-1 (Fig. 4D).

Gross morphology showed that all constructs cultured in either chondrogenic (Fig. 5) or osteogenic (Fig. 6) medium retained their hemispherical architecture throughout the 28-day differentiation period. In basal chondrogenic medium without rhTGF-β3, NT and eGFP-transduced constructs produced little matrix, and the observed staining came from remnants of the CDM hemispheres (Fig. 5). Both TGF-β3-transduced constructs and NT constructs fed rhTGF-β3 synthesized robust cartilaginous matrix that stained positive for GAG and type II collagen and completely filled in the pores of CDM hemispheres (Fig. 5). However even in the presence of rhTGF-β3, rhIL-1 treatment abolished matrix production in NT constructs (Fig. 5). In contrast, IL-1Ra-transduced constructs generated copious amounts of cartilaginous matrix that completely filled CDM hemispheres in the presence (Fig. 5) or absence (Supplemental Fig. 5) of rhIL-1. Neither type I nor type X collagen was observed in any of the chondrogenic constructs at low (Fig. 5) or high magnification (Supplemental Fig. 6). In all constructs cultured in osteogenic medium, a majority of the GAG and type II collagen staining came from the CDM in the hemispheres (Fig. 6). Newly synthesized matrix did not stain positive type X collagen (Fig. 6), and only a minimal amount of type I collagen was found in eGFP-transduced constructs (Fig. 6 & Supplemental Fig. 7).

**Figure 5:**
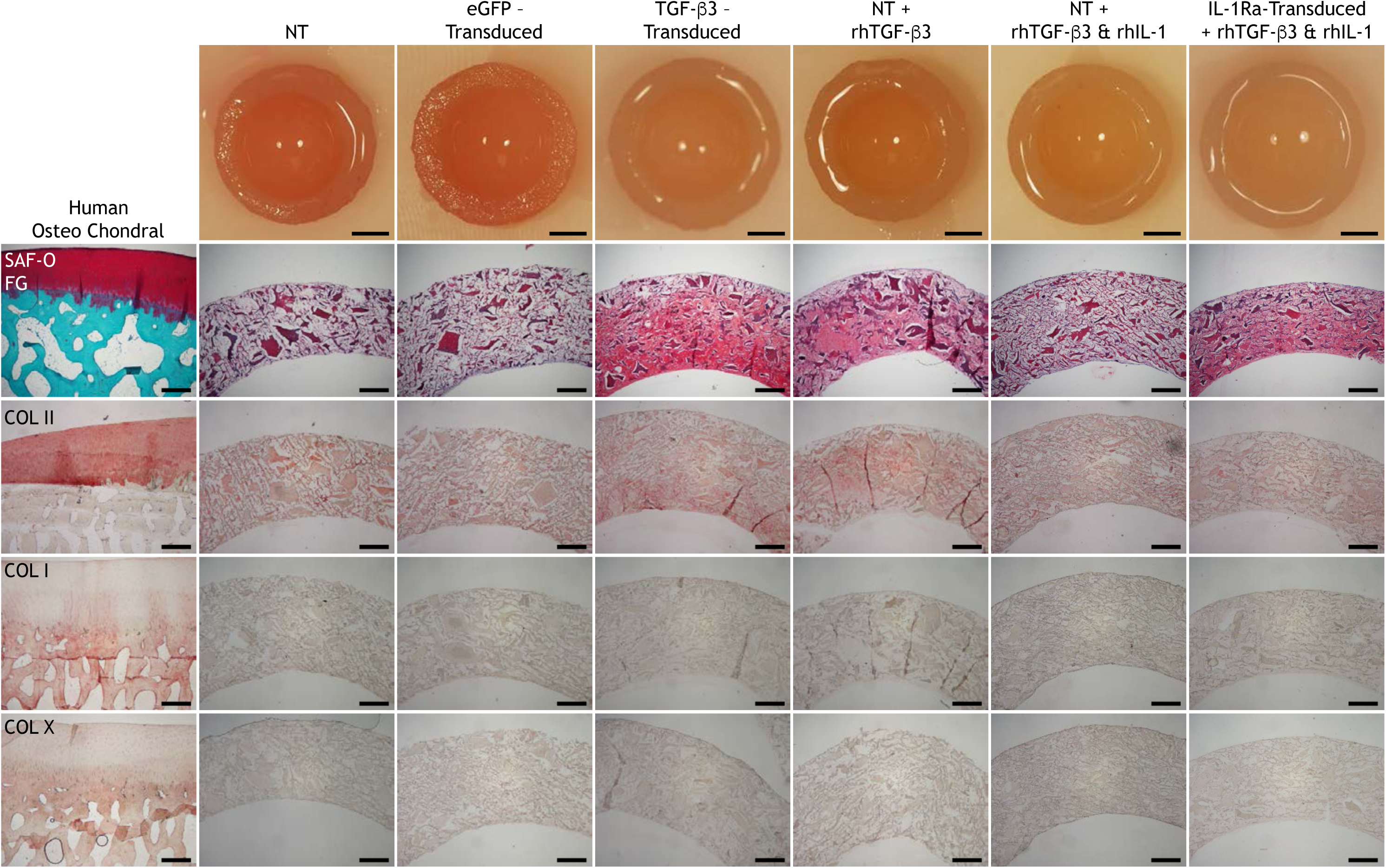
Gross morphology; histology (Safranin-O and Fast Green staining); immunohistochemistry for type II collagen (COL II); type I collagen (COL I); type X collagen (COL X) on human osteochondral sections and non-transduced (NT), eGFP-transduced, or TGF-β3-transduced constructs cultured in basal chondrogenic media (without exogenous rhTGF-β3), NT constructs fed exogenous rhTGF-β3 with and without rhIL-1, and IL-1Ra-transduced constructs fed exogenous rhTGF-β3 and rhIL-1 after 28 days of culture in chondrogenic media. All images are transverse sections, showing full-thickness of each construct. Aminoethyl carbazone (AEC) produces red color. Gross picture scale: 2mm. Histology scale: 500μm.

**Figure 6:**
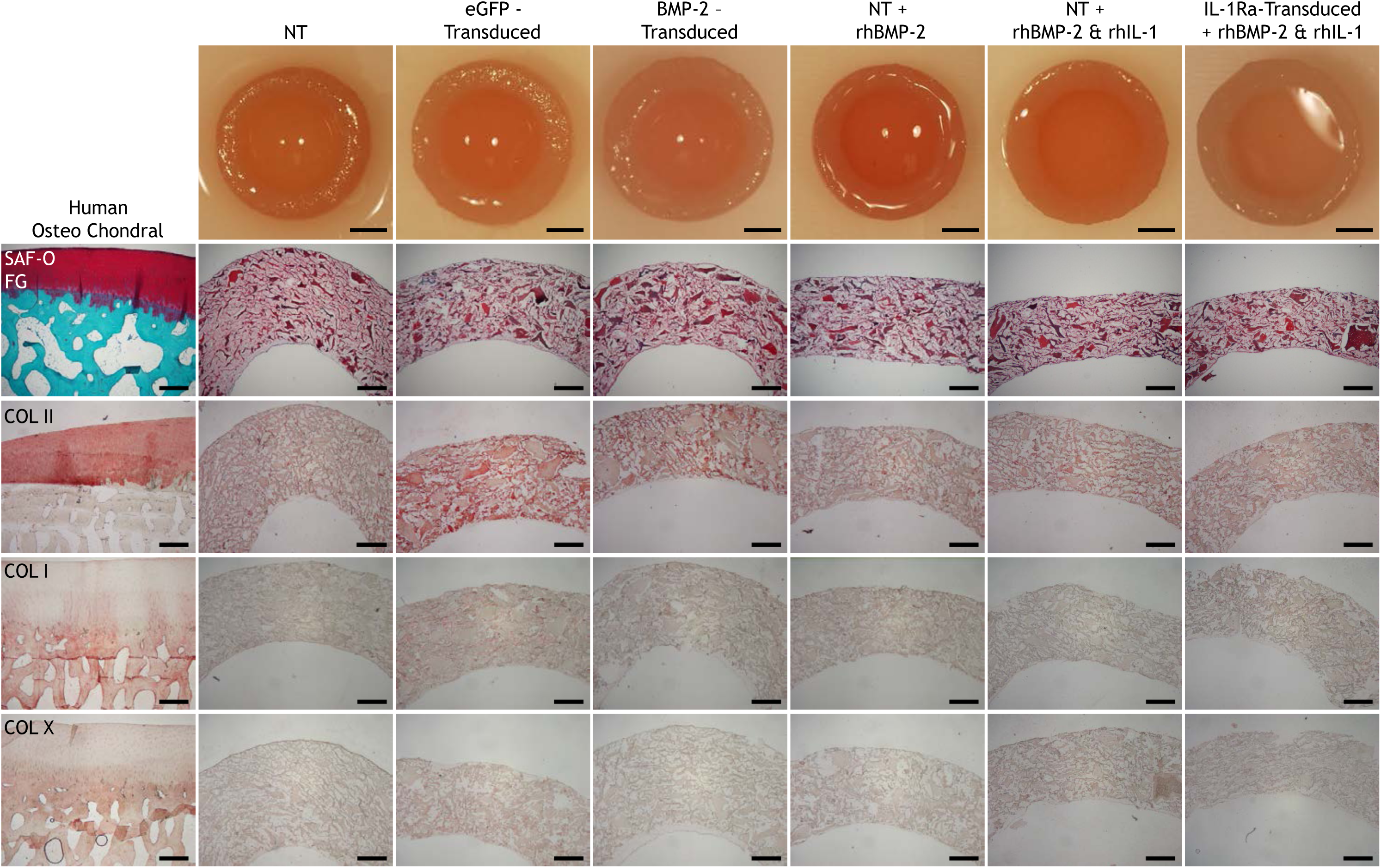
Gross morphology; histology (Safranin-O and Fast Green staining); immunohistochemistry for type II collagen (COL II); type I collagen (COL I); type X collagen (COL X) on human osteochondral sections and non-transduced (NT), eGFP-transduced, or BMP-2-transduced constructs cultured in basal osteogenic media (without exogenous rhBMP-2), and NT constructs fed exogenous rhBMP-2 with and without rhIL-1, and IL-1Ra-transduced constructs fed exogenous rhBMP-2 and rhIL-1 after 28 days of culture in osteogenic media. All images are transverse sections, showing full-thickness of each construct. Aminoethyl carbazone (AEC) produces red color. Gross picture scale: 2mm. Histology scale: 500μm.

Micro-CT revealed that mineralization only occurred in constructs cultured in osteogenic medium (Fig. 7), not in constructs cultured in chondrogenic medium (data not shown). After 28 days of differentiation, both BMP-2-transduced constructs and NT constructs fed rhBMP-2 produced significantly larger volumes of mineral compared to NT and eGFP-transduced constructs cultured in basal osteogenic medium (Fig. 7A). Treatment with rhIL-1 did not have any effect on the mineral volume of NT, eGFP-transduced, or IL-1Ra-transduced constructs (Fig. 7B). For all constructs, the calcium contents measured in the decalcification solutions followed the same trends as the micro-CT data (Fig. 7C & **7D**). While BMP-2 expression and rhBMP-2 supplementation yielded the same quantity of mineralization (Fig. 7A & **7C**), the distribution of mineral deposition was dramatically different (Fig. 7E). In BMP-2-transduced constructs, mineral was evenly distributed and the ratio of mineral produced in the bottom half of constructs to the top half (bottom/top) was 1.00 ± 0.09, whereas in constructs fed rhBMP-2 the mineral was preferentially deposited in the top half of constructs yielding a ratio of 0.68 ± 0.05 (Fig. 7E). This difference in mineral distribution was visually apparent (Supplemental Fig. 8). A representative image showed that newly synthesized mineral was found throughout CDM constructs (Fig. 7F, 7G, 7H).

**Figure 7:**
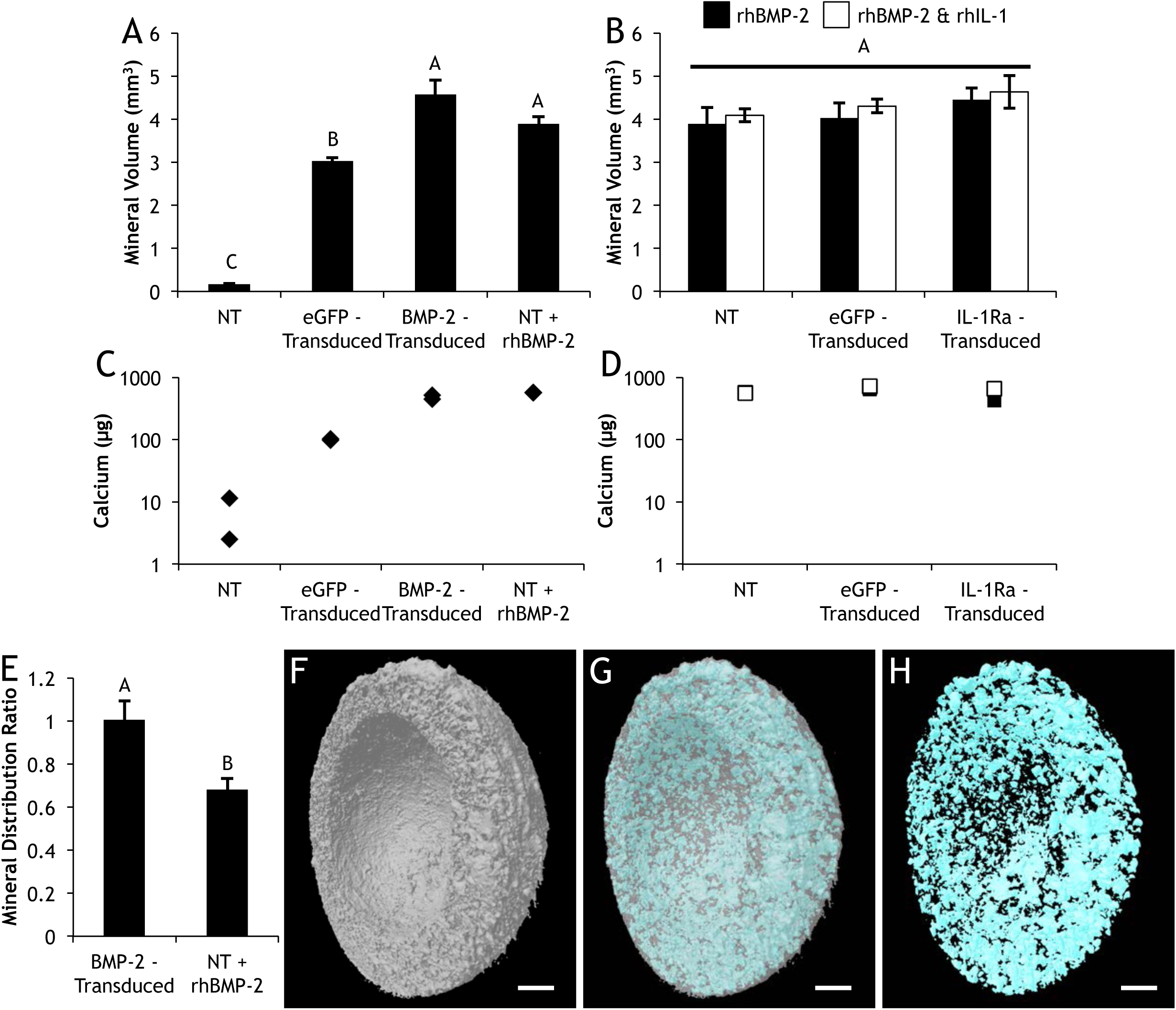
Day 28 (A&B) total volume of mineralization and (C&D) calcium content in (A&C) non-transduced (NT), eGFP-transduced, or BMP-2-transduced constructs cultured in basal osteogenic media (without exogenous rhBMP-2), and NT constructs fed exogenous rhBMP-2; (B&D) NT, eGFP-transduced, and IL-1Ra-transduced constructs cultured in osteogenic media containing rhBMP-2 with and without rhIL-1. (A&B) Bars represent mean ± SEM (n=6). Groups not sharing same letter have p-values < 0.05. (C&D) n=2. (E) Ratio of mineral distribution between the bottom and top of BMP-2-transduced constructs or NT constructs fed exogenous rhBMP-2 (bars represent mean ± SEM, n=6). Representative μCT images of constructs showing (F) CDM, (G) merged, (H) mineral content. Scale: 1mm. There was no detectable mineralization or calcium content in any of the chondrogenic samples.

TGF-β3-transduced hemispherical shells possessed cavities with the exact dimensions of BMP-2-transduced hemisphere cores, and the concentric hemispheres were combined after 22 days of respective chondrogenic or osteogenic culture to form osteochondral constructs (Fig. 8A). Control constructs were transduced with eGFP on both hemispherical shells and hemisphere inserts. Gross morphology demonstrated that osteochondral constructs maintained their hemispherical shape throughout the 50-day differentiation culture (Fig. 8B). At Day 50, TGF-β3 + BMP-2-transduced constructs were more opaque, indicative of new matrix deposition, compared to eGFP-transduced controls (Fig. 8B). Confocal images of eGFP-transduced constructs revealed that both outer hemispherical shells and inner hemisphere cores retained eGFP+ cells a full 56 days after scaffold-mediated lentiviral transduction (Fig. 8C). With administration of 1 μg/mL dox, TGF-β3 + BMP-2-transduced constructs secreted detectable levels of both proteins throughout the 50-day differentiation culture, while both proteins were undetectable in the absence of dox (Fig. 8D). The decrease in protein detected after combining the two-part constructs was likely due to a decrease in surface area exposed to the culture media thereby entrapping growth factors within the constructs (Fig. 8D). TGF-β3 + BMP-2-transduced constructs produced robust quantities of newly synthesized matrix that completely filled in pores of outer hemispherical shells, inner hemisphere cores, and the interface between the two-part constructs (Fig. 9). In growth factor-transduced constructs, newly deposited matrix exhibited a spatially distinct composition; both GAG and type II collagen were primarily localized to TGF-β3-transduced hemispherical shells, whereas type I collagen was found exclusively in the BMP-2-transduced hemisphere cores (Fig. 9). Interestingly, the matrix deposited in the interface, which fused the two-part constructs together, stained for GAG, type II, and type I collagen (Fig. 9). Constructs expressing eGFP produced negligible amounts of matrix and remained as two separate pieces that did not integrate (Fig. 9). While there was no difference in total collagen content (Fig. 10A), the TGF-β3 and BMP-2 expressing osteochondral constructs produced significantly higher GAG (Fig. 10B), DNA (Fig. 10C), and mineral contents (Fig. 10D) compared to eGFP expressing constructs, and mineral deposition co-localized with BMP-2 expression in the inner hemisphere core (Fig. 10E, 10F, 10G).

**Figure 8:**
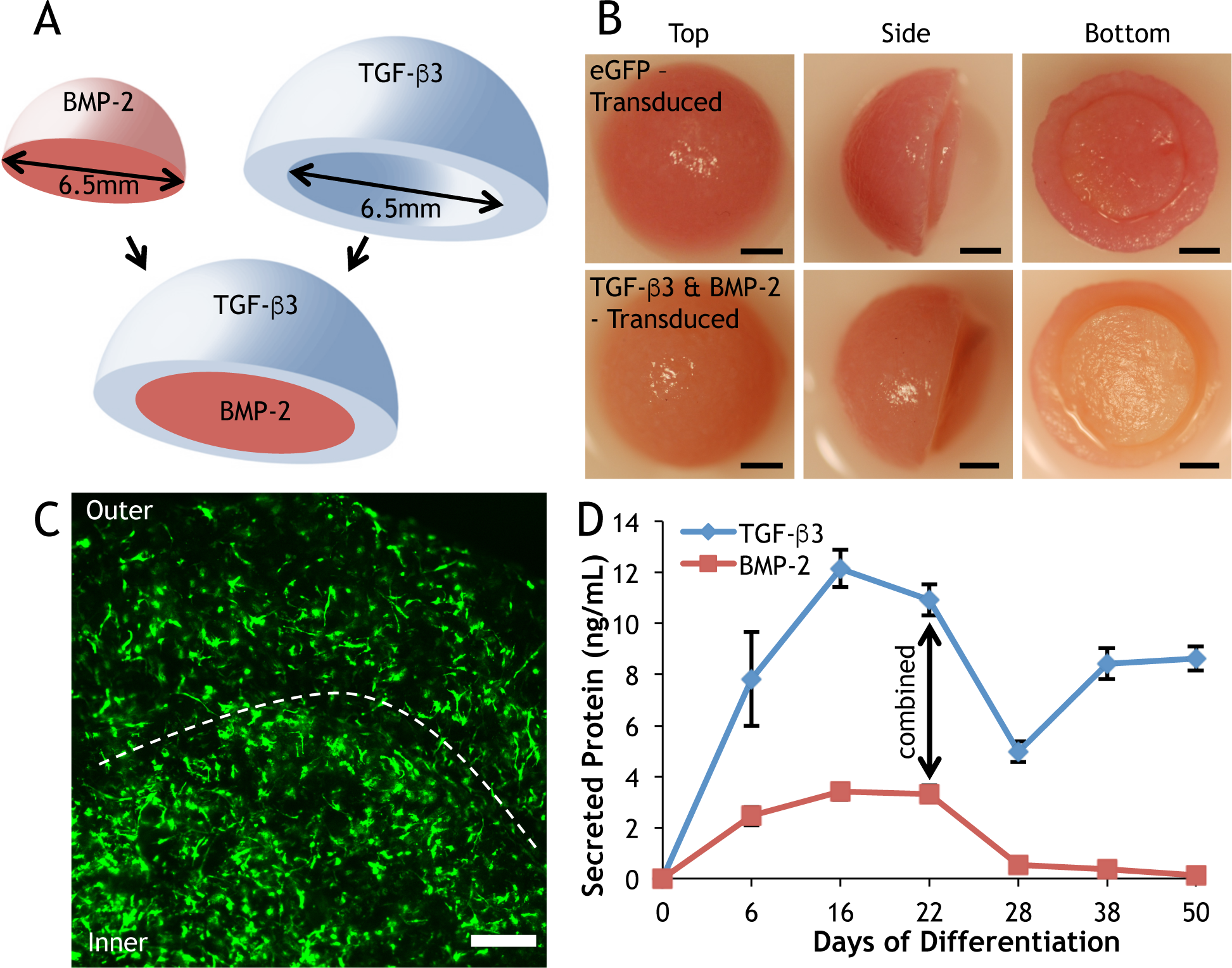
Osteochondral constructs after 22 days of respective osteogenic or chondrogenic differentiation followed by 28 days of co-culture. (A) Illustration of chondrogenic outer hemispherical shell with immobilized TGF-P3 lentivirus and osteogenic inner hemisphere core with immobilized BMP-2 lentivirus. (B) Day 50 gross morphology of the top, side, and bottom of eGFP-transduced or TGF-P3 + BMP-2-transduced constructs. Scale: 2 mm. (C) Confocal image of eGFP-transduced MSCs in the inner hemisphere core and outer hemispherical shell at Day 50 of differentiation. Interface denoted by dotted line. Scale: 500um. (D) Growth factor secretion of TGF-P3 + BMP-2-transduced osteochondral constructs combined at Day 22 (mean ± SEM, n=8).

**Figure 9:**
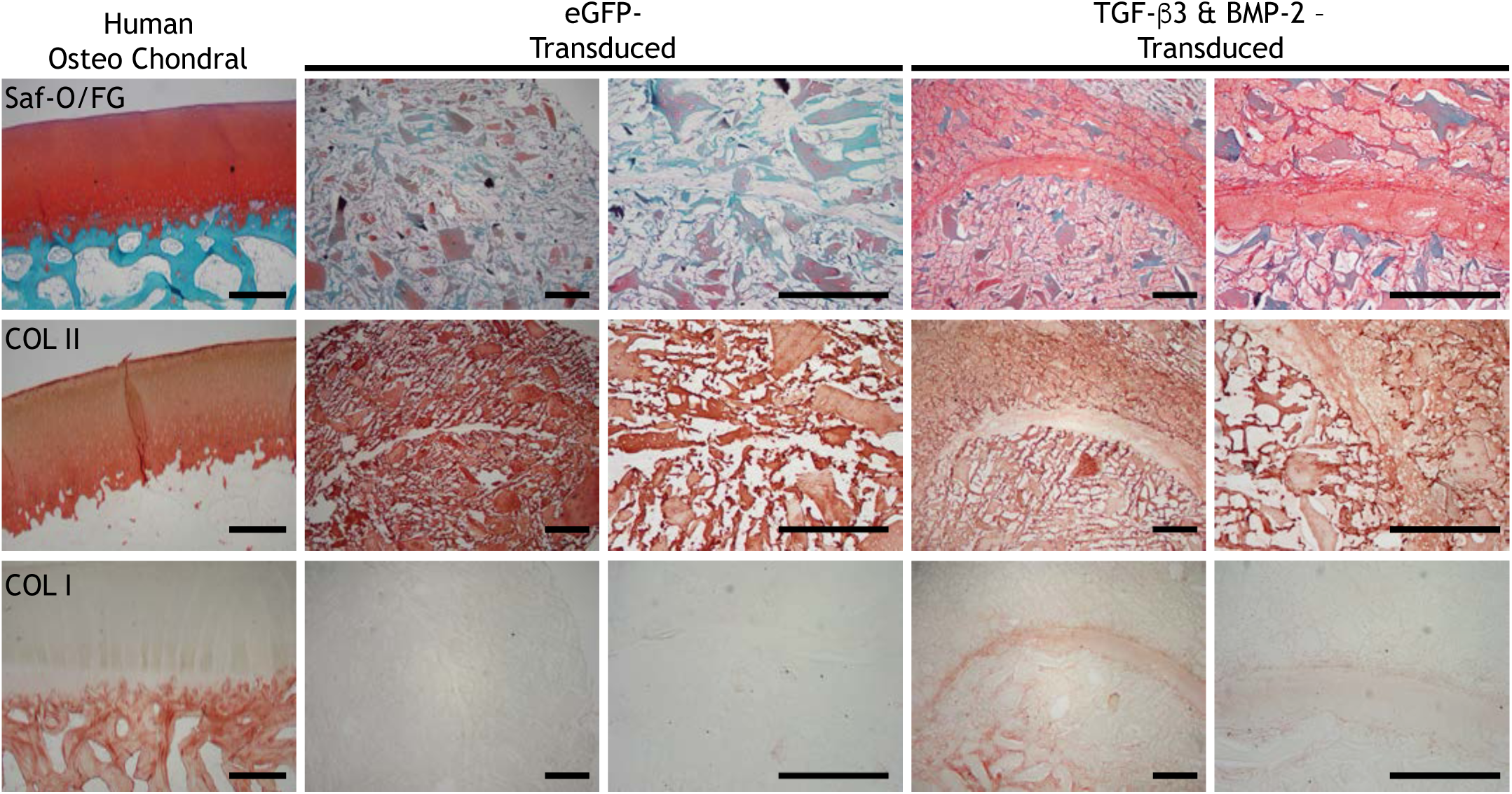
Histology (Safranin-O and Fast Green staining); immunohistochemistry for type II collagen (COL II); type I collagen (COL I); on human osteochondral sections and eGFP-transduced or TGF-P3 + BMP-2-transduced constructs after 50 days of differentiation. For CDM constructs, first column taken at 4x, second column taken at 10x. All images are transverse sections, showing full-thickness of each construct. Aminoethyl carbazone (AEC) produces red color. Scale: 500um.

## Discussion

The results of this study demonstrate that both tissue remodeling and osteochondral matrix production can be spatially and temporally regulated within anatomically-shaped CDM constructs via scaffold-mediated lentiviral gene delivery of dox-inducible anti-inflammatory cytokines and tissue-specific growth factors. Scaffold-mediated lentiviral gene delivery provided highly efficient, site-specific transduction of all transgenes of interest, whereas the use of a dox-inducible promoter afforded inducible and tunable control over protein secretion. The controlled induction of IL-1Ra in MSCs offered a novel approach for exogenously regulating the cell-mediated degradation of CDM constructs under inflammatory conditions without modifying the intrinsic properties of CDM via chemical crosslinking. In the controlled induction of MSC differentiation, we found that secretion of TGF-β3 and BMP-2 were equally as effective at driving respective chondrogenic and osteogenic differentiation as exogenous supplementation of the same growth factors, but possessed the added benefit of yielding more uniform matrix deposition compared to exogenous growth factor supplementation. The bi-layered nature of the osteochondral constructs prevented the interference of antagonistic growth factor signaling and generated robust, tissue-specific differentiation. Furthermore, the ability of CDM to undergo cellular remodeling served as an effective tool for promoting integration of the two parts without the need for external fixatives or adhesives. These findings show that decellularized CDM can provide a single, homogenous biomaterial scaffold for fabricating anatomically-shaped osteochondral constructs, and that scaffold-mediated lentiviral gene delivery can confer spatial and temporal regulation over both scaffold remodeling and site-specific osteochondral differentiation.

We found that using PLL to immobilize lentivirus to the scaffolds did not affect the biochemical composition of CDM hemispheres (Fig. 4 **Caption**) or influence MSC attachment (Fig. 1B) or viability (Supplemental Fig. 3). The cationic nature of PLL neutralizes the negatively charged viral vectors and facilitates noncovalent immobilization onto synthetic [55, 62, 63] and collagen [67] scaffolds. While these studies demonstrated that PLL treatment was well tolerated by cells [55, 62, 63, 67], a critical objective of the current investigation was to verify that lentivirus immobilization did not negatively impact the native structure or composition of CDM, which in turn influence cell attachment and viability. Minimizing alterations to CDM is of the utmost importance as processing techniques including excessive chemical crosslinking [47, 68], pepsin digestion [34, 69], encapsulation in PLGA microspheres [23, 70], incorporation with carbon nanotubes [71], or functionalization with methacrylate groups [17, 72] can inhibit cell attachment [47, 68, 71] and compromise the ability of CDM to support chondrogenesis [47, 68–70] or endochondral ossification [17]. In the current study, there was no difference in the baseline biochemical composition between NT and eGFP-transduced constructs (reported in the caption of Fig. 4). These values are in agreement with previously reported biochemical values for CDM hemispheres [49]. Additionally, SEM illustrated that MSCs readily adhered to the collagen fibrils present in CDM hemispheres containing immobilized lentivirus (Fig. 1B). Moreover, the MSCs adopted a cell morphology (Fig. 1B) traditionally observed in non-PLL-coated CDM scaffolds without viral vectors [73, 74], which indicates that the PLL treatment did not negatively affect cell interactions with CDM.

Scaffold-mediated lentiviral gene delivery to seeded MSCs allowed for inducible protein synthesis for the entire culture period (Fig. 2 & Supplemental Fig. 4). Various components within CDM are capable of binding growth factors, and this ability motivated the use of CDM as a growth factor-eluting reservoir both *in vitro* [75–78] and *in vivo* [79–81]. However, these approaches may be limited by a burst release at the beginning of the culture period [75, 77–80], the need for supraphysiological concentrations in order to stimulate chondrogenesis [75–77, 79], and relatively short timelines (on the order of days) for growth factor availability [77–81]. These challenges are further compounded by the very short *in vivo* half-lives of the active forms of TGF-β3 [82], BMP-2 [83], and IL-1Ra [84] which has motivated the use of scaffold-mediated gene delivery as a strategy for spatially and temporally regulating growth factor [56] and anti-inflammatory cytokine [55, 63] presentation. While scaffold-mediated, non-viral [57] and adeno-associated viral [59–61] gene delivery have been extensively explored for both cartilage and bone regeneration, low transfection efficiency and transient expression limit the efficacy and applicability of those approaches. In one study investigating TGF-β3 and BMP-2 delivery for osteochondral tissue engineering, cells were incubated with plasmids prior to encapsulation within alginate hydrogels in order to increase the transfection efficiency, necessitating a multi-step transfection process [15]. Other non-viral approaches exploited various biomaterial properties to temporally control gene delivery; however, the biomaterials solely served to prolong plasmid uptake to increase expression duration as the plasmid is diluted when cells proliferate [3, 4, 8]. In contrast, lentivirus can transduce both dividing and non-dividing cells with high transduction efficiency and integrates transgenes into the host genome, allowing for indefinite expression [85]. In CDM hemispheres, scaffold-mediated lentiviral gene delivery yielded a transduction efficiency in MSCs of 57 ± 5%, which was much higher than the 20% transfection efficiency observed in non-viral plasmid delivery to MSCs in alginate hydrogels [15]. However, this transduction efficiency was lower than the 70–85% transduction efficiencies observed with scaffold-mediated lentiviral gene delivery on synthetic PCL scaffolds [55, 62, 63]. These discrepancies could be due to differences in how cells respond to natural versus synthetic materials, as previous studies have revealed that substrate composition [86], topography [86], stiffness [87], and ligand presentation [87] influence cell morphology, motility, and transfectability [86, 87]. Despite the lower transduction efficiency observed, the concentrations of secreted growth factors (Fig. 2 A & B) and anti-inflammatory cytokines (Fig. 2 C) in this study were similar to the concentrations of TGF-β3 [62] and IL-1Ra [55] generated by MSCs in PCL constructs, and were an order of magnitude higher than the concentrations of TGF-β3 and BMP-2 synthesized by non-viral plasmid delivery [8, 15]. While previous studies have utilized constitutive promoters to drive growth factor expression [4, 8, 62], the persistent expression of growth factors induced a hypertrophic phenotype in MSCs [61], and may be undesirable for *in vivo* application [61]. Using biomaterials to prolong non-viral gene delivery does not afford the type of temporal regulation needed to address this issue; rather, dox-inducible promoters [88, 89] have been shown to control both the magnitude and duration of transgene expression, and enable intermittent activation of transgene expression [55]. This degree of temporal control is critical for *in vivo* success as stem cells exhibit a donor-dependent response to growth factor release from CDM scaffolds [77], and the dox-inducible promoters allow for growth factor expression to be tailored in an exogenously controlled manner.

IL-1Ra secretion (Fig. 2C) inhibited rhIL-1-mediated MMP activity (Fig. 3) and prevented enzymatic degradation of CDM constructs (Fig. 4B & Fig. 4D). Modulating the inflammatory responses towards tissue-derived scaffolds is critical for promoting successful outcomes *in vivo* [32], particularly in conditions of injury or disease. In the current study, CDM was decellularized prior to hemisphere fabrication, which significantly mitigates host inflammatory responses [90]. However, removal of antigenic components alone may not be sufficient, and recellularization with MSCs has been shown to play an active role in regulating the inflammatory environment [91, 92]. One of the mechanisms by which MSCs exert their immunomodulatory effects is through the secretion of IL-1Ra, which encourages macrophage polarization towards a M2 phenotype *in vitro* and inhibits B cell differentiation *in vivo* [53]. Therefore, scaffold-mediated lentiviral gene delivery of dox-inducible IL-1Ra provided tunable, exogenous control over the native immunomodulatory properties of MSCs. In the presence of IL-1, previous studies have demonstrated that viral transduction of IL-1Ra preserves the chondrogenic capacity of MSCs and inhibits the catabolic effects of IL-1 in pellets [93] and on synthetic scaffolds [55, 63]. While these studies elucidated the capacity of IL-1Ra expression to prevent the degradation of cell-synthesized neomatrix, the current study extends this concept by investigating whether IL-1Ra secretion could also prevent degradation of cell-labile CDM hemispheres. Following scaffold-mediated lentiviral gene delivery, dox treatment resulted in IL-1Ra accumulation above 100 ng/mL (Fig. 2C), which was 1000-fold higher than the physiologic concentration of rhIL-1α (0.1 ng/mL) administered and was sufficient to suppress MMP upregulation throughout the entire study (Fig. 3). These results are consistent with the IL-1Ra concentration produced in pellets [93] and synthetic scaffolds [55] virally transduced to express IL-1Ra. An important finding of the present study was the elucidation that modulation of MMP activity (Fig. 3) had a profound effect on the collagen content of CDM hemispheres (Fig. 4B & Fig. 4D). At physiologic concentrations of rhIL-1α, NT and eGFP-transduced CDM constructs possessed upregulated MMP activity (Fig. 3), which corresponded to respectively lower collagen contents in both chondrogenic (Fig. 4B) and osteogenic (Fig. 4D) media. These results are in agreement with previous findings that revealed IL-1α increases MMP expression leading to subsequent collagen degradation [94]. In contrast, IL-1Ra-transduced CDM constructs did not exhibit any increase in MMP activity (Fig. 3) and retained the native collagen content throughout the entire 28-day culture period (Fig. 4B & Fig. 4D). IL-1Ra secretion did not completely abolish MMP activity, but rather maintained MMP activity in its basal state in MSCs in the absence of IL-1 (Fig. 3). This basal level of MMP activity may be critical to preserve because MMP expression has been shown to influence both chondrogenic and osteogenic differentiation of MSCs [95], suggesting that the ability of cells to remodel their environment plays a prominent role in regulating differentiation. In this regard, our findings show proof-of-concept that controlling scaffold degradation through modulating the response to inflammation can reduce degradation while preserving the inherent capacity of CDM to undergo cellular remodeling. On the other hand, chemically crosslinking scaffolds to inhibit enzymatic degradation [45, 46] not only leads to fibrotic encapsulation [32], but also prevents the generation of degradation products [96]. The degradation products of tissue-derived scaffolds have not only been shown to promote a constructive remodeling M2 macrophage phenotype [30, 31] but also to stimulate chondrogenic differentiation [33, 34]. Importantly, neither dehydrothermal treatment nor lentiviral immobilization affected the ability of CDM hemispheres to degrade as eGFP-transduced constructs still degraded in the presence of rhIL-1 in both chondrogenic (Fig. 4B) and osteogenic (Fig. 4D) media. These results show that scaffold-mediated lentiviral gene delivery in and of itself does not affect the degradation of CDM hemispheres, but rather the secretion of dox-inducible IL-1Ra was the mediating factor for regulating scaffold degradation. Therefore, the combination of inducible gene delivery and physical crosslinking of the scaffold [47] provided a unique means of altering the degradation rate of CDM hemispheres while preserving the capacity of CDM to undergo cellular remodeling in an inflammatory environment.

Biochemical composition (Fig. 4), histological and immunohistochemical staining (Fig. 5 & Fig. 6), and mineralization data (Fig. 7) collectively demonstrated that scaffold-mediated lentiviral gene delivery stimulated tissue-specific matrix production in CDM constructs. These results agree with both the typical chondrogenic response observed in CDM hemispheres [49] and the chondrogenic differentiation stimulated by scaffold-mediated lentiviral gene delivery of TGF-β3 [62]. In contrast, either exogenous supplementation or viral expression of BMP-2 produced substantial mineralization (Fig. 7) but yielded negligible GAG synthesis (Fig. 4C & Fig. 6). Interestingly, BMP-2-transduced constructs (~10 ng/mL BMP-2) and constructs fed rhBMP-2 (100 ng/mL) deposited equivalent amounts of mineralization (Fig. 7A & **7C**) despite an order of magnitude difference in concentrations of BMP-2 driving differentiation. Previous studies with growth factor-absorbed CDM scaffolds demonstrated a similar finding where lower concentrations of growth factors were capable of driving differentiation potentially due to increased bioavailability and accessibility to seeded cells [78, 80]. Alternatively, the concentration of BMP-2 secreted by transduced constructs could be much higher than measured and be sequestered inside CDM hemispheres by binding to various components within the CDM [77]. The ability of CDM to bind and sequester growth factors could also produce the heterogeneity of mineralization observed in the constructs fed rhBMP-2 (Fig. 7E & Supplemental Fig. 8), as diffusion gradients of supplemented growth factors produce heterogeneous matrix deposition within tissue-engineered constructs [97]. Therefore, scaffold-mediated lentiviral gene delivery not only produced equivalent quantities of respective cartilaginous and osseous matrix but also provided the added benefit of generating homogenous mineral distribution. Interestingly, there was no effect of rhIL-1α treatment or IL-1Ra expression on mineral content (Fig. 7B & **7D**). While IL-1β has been shown to enhance endochondral ossification of MSCs [98, 99], these studies did not include BMP-2, which might saturate the capacity of MSCs to deposit calcium. Alternatively, studies investigating the effects of IL-1β within osteochondral constructs have demonstrated that MSCs respond differently to IL-1β depending on their differentiation state [11]; therefore, the ability to spatially and temporally regulate IL-1Ra secretion could play an important role in controlling inflammatory signaling within disparate tissue types as well as modulating the inflammatory signaling across the osteochondral interface.

While these results demonstrated that scaffold-mediated lentiviral gene delivery of TGF-β3 and BMP-2 could drive respective chondrogenic and osteogenic differentiation in isolated CDM hemispheres, the final aim utilized scaffold-mediated transduction to spatially pattern tissue differentiation within concentric CDM hemispherical shells and hemisphere cores in order to fabricate anatomically-shaped, osteochondral constructs. Indeed, spatially regulating tissue-specific growth factor secretion yielded hierarchically organized matrix deposition phenotypic of cartilage and bone. GAG and type II collagen staining (Fig. 9) colocalized with outer TGF-β3-transuced hemispherical shells (Fig. 8A), while type I collagen staining (Fig. 9) and mineralization (Fig. 10) were restricted to the inner BMP-2-transduced hemisphere cores (Fig. 8A). In contrast with previous studies that incorporated CDM into biphasic constructs with the subchondral bone region composed of demineralized bone matrix [23, 36, 37], hydroxyapatite [100], calcium phosphate [44], or synthetic polymers [38], the current study generated both cartilaginous and osseous phases from a single, compositionally homogenous CDM substrate. Furthermore, the current study demonstrated that lentiviral transduction produced robust growth factor synthesis for the entire 50-day culture period, which did not peak until Day 16 (Fig. 8D). These results contrast with studies that spatially stratified plasmids encoding TGF-β1 [4] or -β3 [8] along with BMP-2 and found that growth factor production peaked at Day 2 [8] and only lasted 14 days [4]. These discrepancies are likely due to the highly efficient lentiviral transduction and stable expression afforded by integration [85] as evidenced by the confluent eGFP expression a full 56 days after scaffold-mediated transduction (Fig. 8C). Generating anatomically-shaped cartilage constructs using woven [63] or 3D-printed [101] PCL, or methacrylated hyaluronic acid scaffolds [102] face the challenge of anchoring to and integrating with the underlying subchondral bone [102], which has motivated engineering anatomically-shaped osteochondral constructs [7, 9, 10]. A major limitation to these approaches is their reliance on non-degradable agarose [7, 9], which could lead to fibrotic encapsulation, or PCL [10], which has been shown to stimulate osseous wall resorption and widen osteochondral defects [6]. In previous reports, anatomically-shaped osteochondral constructs were engineered from two-part scaffolds consisting of alginate and agarose or scaffold-free culture that used fibrin glue to stabilize the interface [9]. In the present study, the concentric CDM hemispheres were able to fuse into a single osteochondral construct without any external fixation required and retained their anatomical conformation throughout the 50-day culture period (Fig. 8B). The osteochondral CDM constructs integrated through matrix deposition and elaboration, which spanned the interface of the concentric hemispheres expressing growth factors (Fig. 9). Additionally, CDM scaffold remodeling could have contributed to this integration via the basal MMP activity observed throughout the study (Fig. 3).

**Figure 10:**
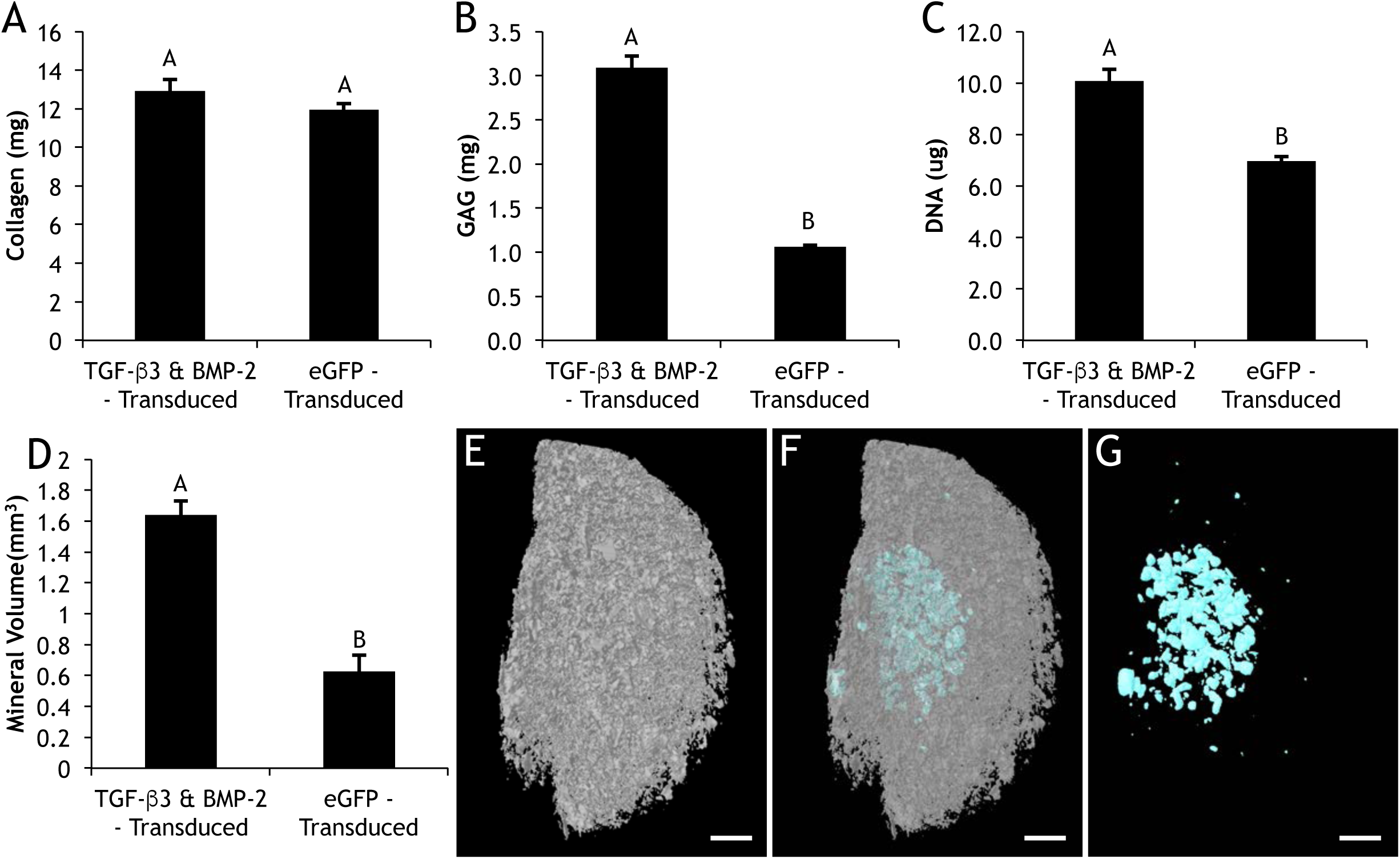
Day 50 (A) collagen; (B) GAG; (C) DNA; (D) total volume of mineralization of eGFP-transduced (n=3) or TGF-P3 + BMP-2-transduced constructs (n=6). Mean ± SEM. Groups not sharing same letter have p-values < 0.05. Representative uCT images of constructs showing (E) CDM, (F) merged, (G) mineral content. Scale: 1mm.

## Conclusions

Scaffold-mediated lentiviral gene delivery of dox-inducible cytokine inhibitors and growth factors enabled spatial and temporal control over both scaffold degradation and osteochondral tissue formation. This system offers a promising strategy for osteochondral repair, and demonstrates the potential for patient-specific tuning of both growth factor secretion and scaffold remodeling. Additionally, the osteochondral constructs provide an *in vitro* model for further investigating the complex interplay of inflammatory signaling within a multi-tissue joint organoid. Therefore, future directions include both the pursuit of clinically-relevant therapies as well as the investigation of the mechanisms of inflammatory signaling across the osteochondral interface.

## Acknowledgments

The authors thank Sara Oswald for providing technical writing support for the manuscript, Matt Joens for assistance in SEM imaging, and Drs. Jonathan Brunger and Charles Gersbach for important discussions and advice in the early stages of the study. This study was supported in part by NIH grants AR50245, AR48852, AG15768, AR48182, AR067467, AR065956, the Nancy Taylor Foundation for Chronic Diseases, the Arthritis Foundation, and the Collaborative Research Center of the AO Foundation, Davos, Switzerland.

---

**Supplemental Figure 1:**
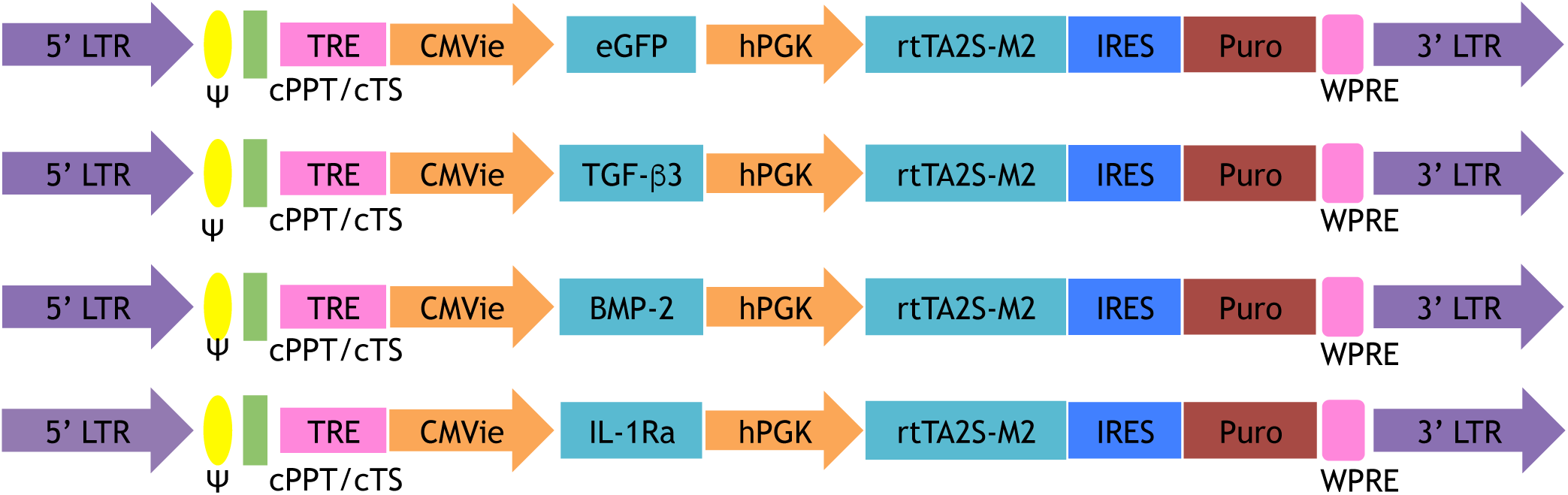
Schematic diagram of dox-inducible lentiviral vectors with eGFP (1st row), TGF-β3 (2nd row), BMP-2 (3rd row), and IL-1Ra (4th row) as the respective genes of interest. Genes of interest are driven by the tetracycline-regulated minimal CMV promoter (TRE-CMVmin). The human phosphoglycerate kinase (hPGK) promoter constitutively drives expression of the tet-responsive transactivator (rtTA2S-M2) and then, following an internal ribosomal entry site (IRES), the puromycin resistance gene (puro) which enables selection of transduced cells. Both vectors contain the 5’ and 3’ long-terminal repeats (LTR), psi packaging signal (Ψ), the central polypurine tract (cPPT), central termination sequence (cTS), and the woodchuck hepatitis virus post-transcriptional regulatory element (WPRE).

**Supplemental Figure 2:**
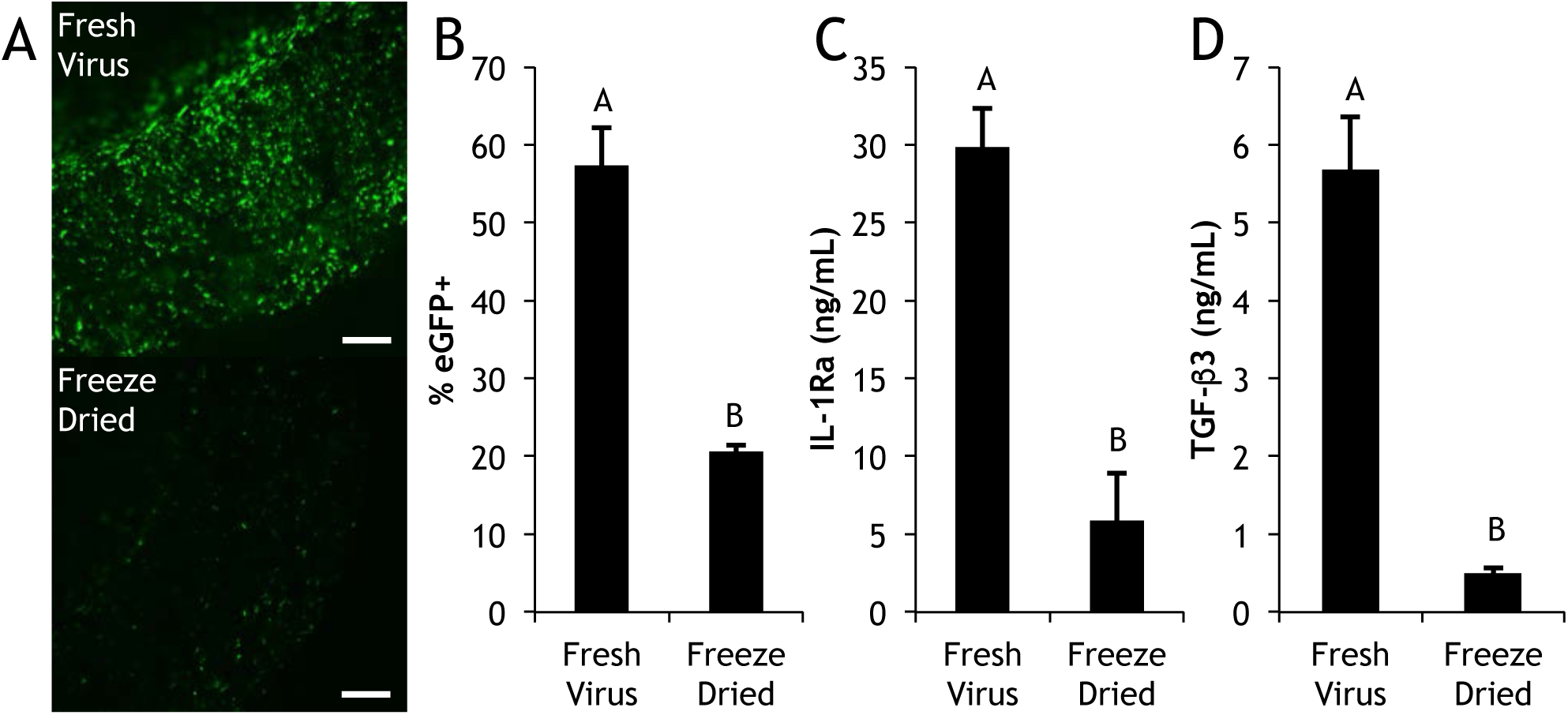
Transduction efficiency of lentivirus immobilized fresh or after being freeze-dried onto CDM hemispheres measured 20 days after biomaterial-mediated transduction. (A) Confocal images of cross-sections of eGFP-transduced constructs. Scale: 500μm. (B) Flow cytometry showing% eGFP+ cells. Protein secretion of (C) IL-1Ra-transduced and (D) TGF-β3-transduced constructs. Mean ± SEM (n=3). Groups not sharing same letter have p-values < 0.05.

**Supplemental Figure 3:**
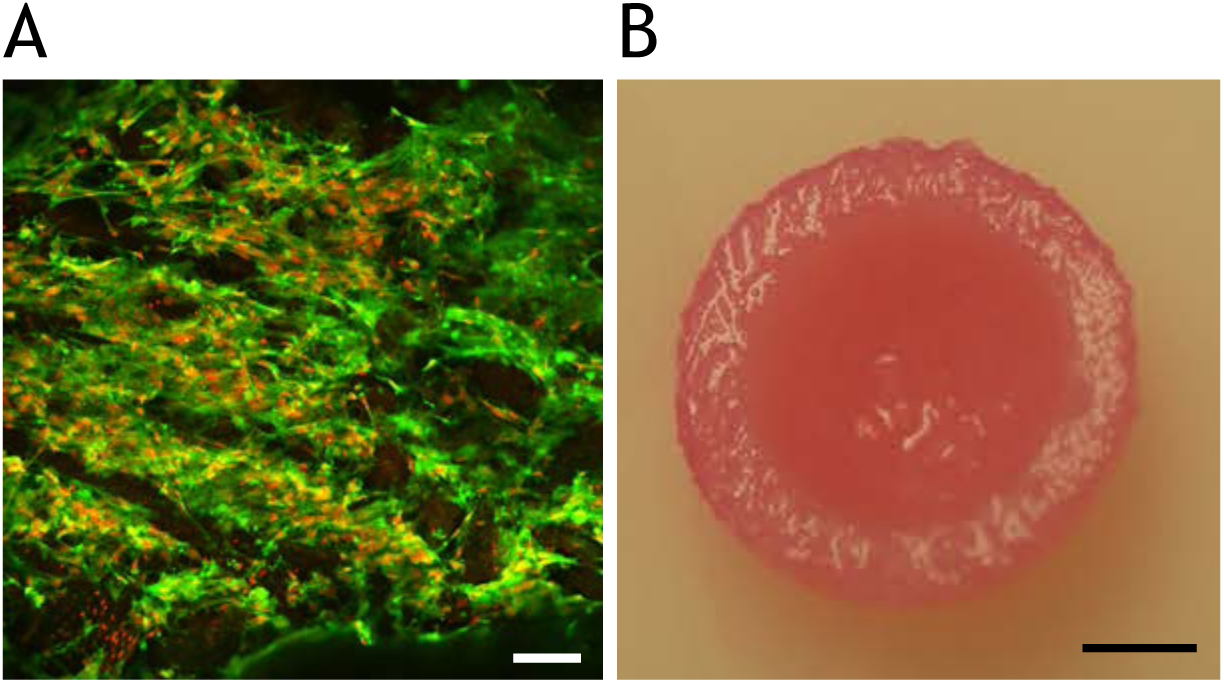
Day 0 (A) confocal image of a non-transduced construct stained for Alexa Phalloidin (f-actin, green) & Propidium Iodide (nuclei, red), scale: 100 **u**m and (B) gross morphology, scale: 2 mm. Confocal image is a transverse section, showing full-thickness of construct.

**Supplemental Figure 4:**
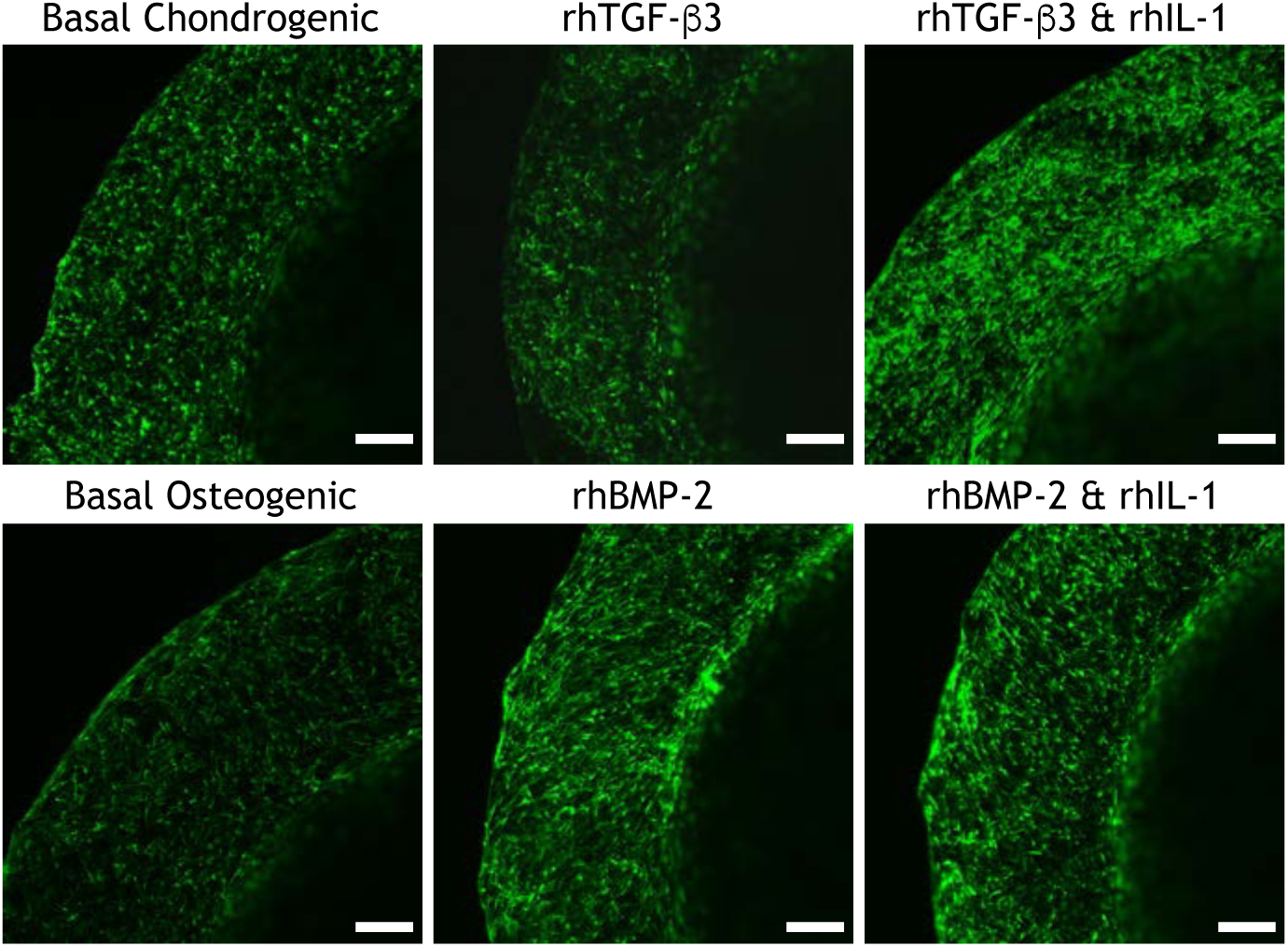
Confocal images of eGFP-transduced constructs after 28 days of culture in chondrogenic (Top) or osteogenic (Bottom) basal media (Left), with exogenous rhTGF-β3 or rhBMP-2 (Middle), and with exogenous rhIL-1 combined with exogenous rhTGF-β3 or rhBMP-2 (Right). All images are transverse sections, showing full-thickness of each construct. Scale: 500 µm.

**Supplemental Figure 5:**
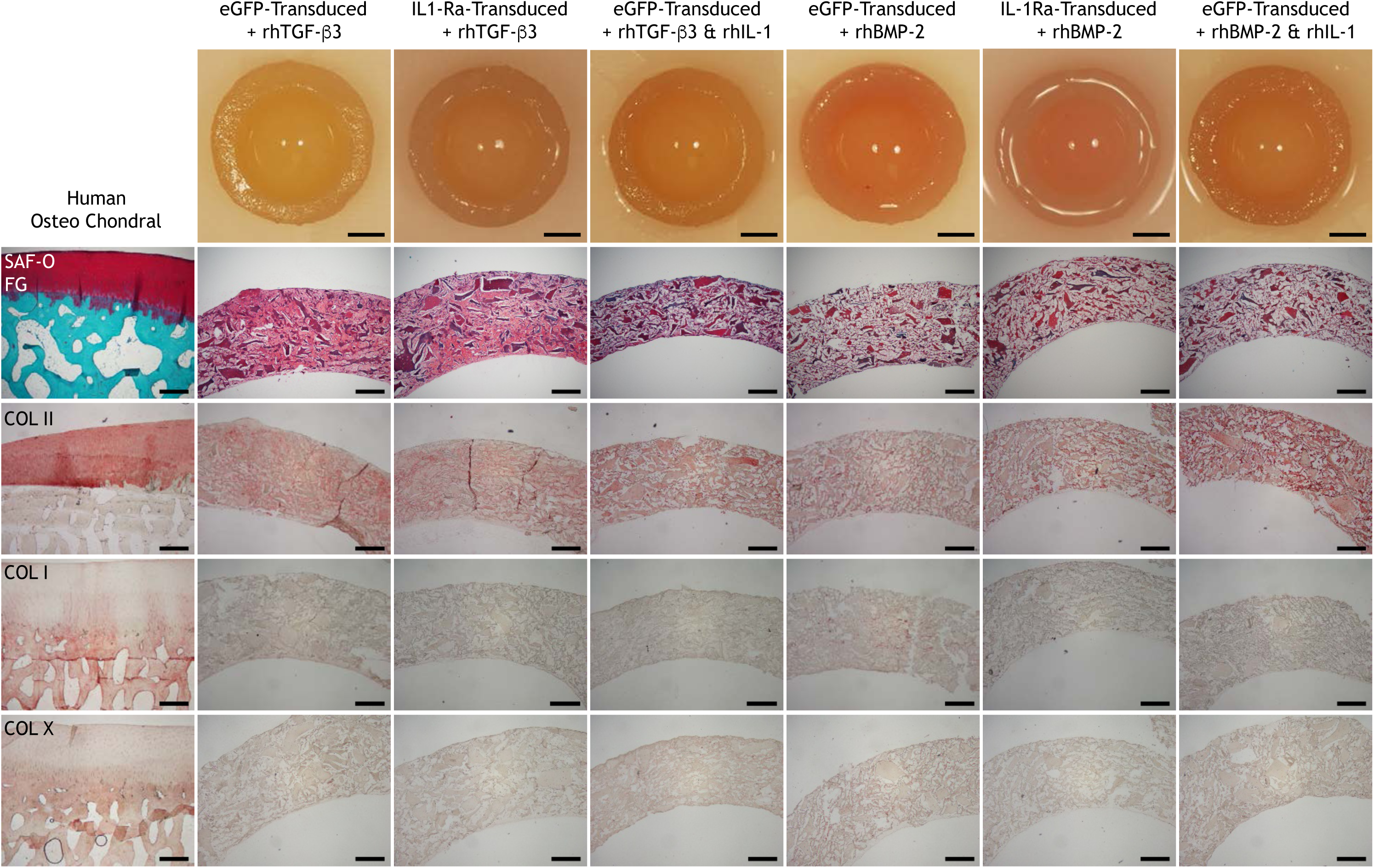
Gross morphology; histology (Safranin-O and Fast Green staining); immunohistochemistry for type II collagen (COL II); type I collagen (COL I); type X collagen (COL X) on human osteochondral sections and eGFP-transduced or IL-1Ra-transduced constructs cultured in chondrogenic media (first three columns) with exogenous rhTGF-β3 or osteogenic media (last three columns) with exogenous rhBMP-2 in the presence and absence of rhIL-1 after 28 days of culture. All images are transverse sections, showing full-thickness of each construct. Aminoethyl carbazone (AEC) produces red color. Gross picture scale: 2mm. Histology scale: 500μm.

**Supplemental Figure 6:**
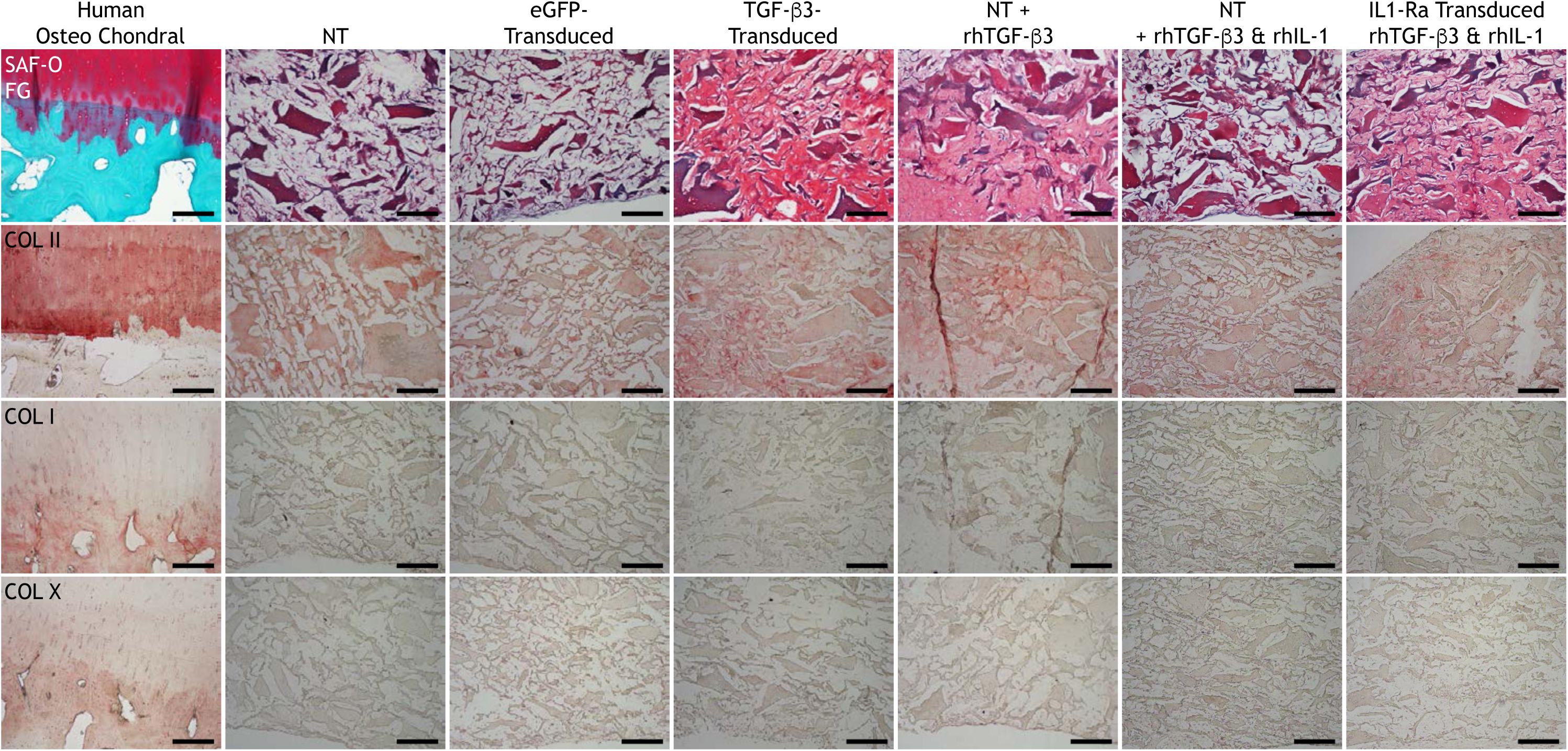
High magnification (10x) histology (Safranin-O and Fast Green staining); immunohistochemistry for type II collagen (COL II); type I collagen (COL I); type X collagen (COL X) on human osteochondral sections and non-transduced (NT), eGFP-transduced, or TGF-β3-transduced constructs cultured in basal chondrogenic media (without exogenous rhTGF-β3), NT constructs fed exogenous rhTGF-β3 with and without rhIL-1, and IL-1Ra-transduced constructs fed exogenous rhTGF-β3 and rhIL-1 after 28 days of culture in chondrogenic media. All images are transverse sections, showing full-thickness of each construct. Aminoethyl carbazone (AEC) produces red color. Histology scale: 250μm.

**Supplemental Figure 7:**
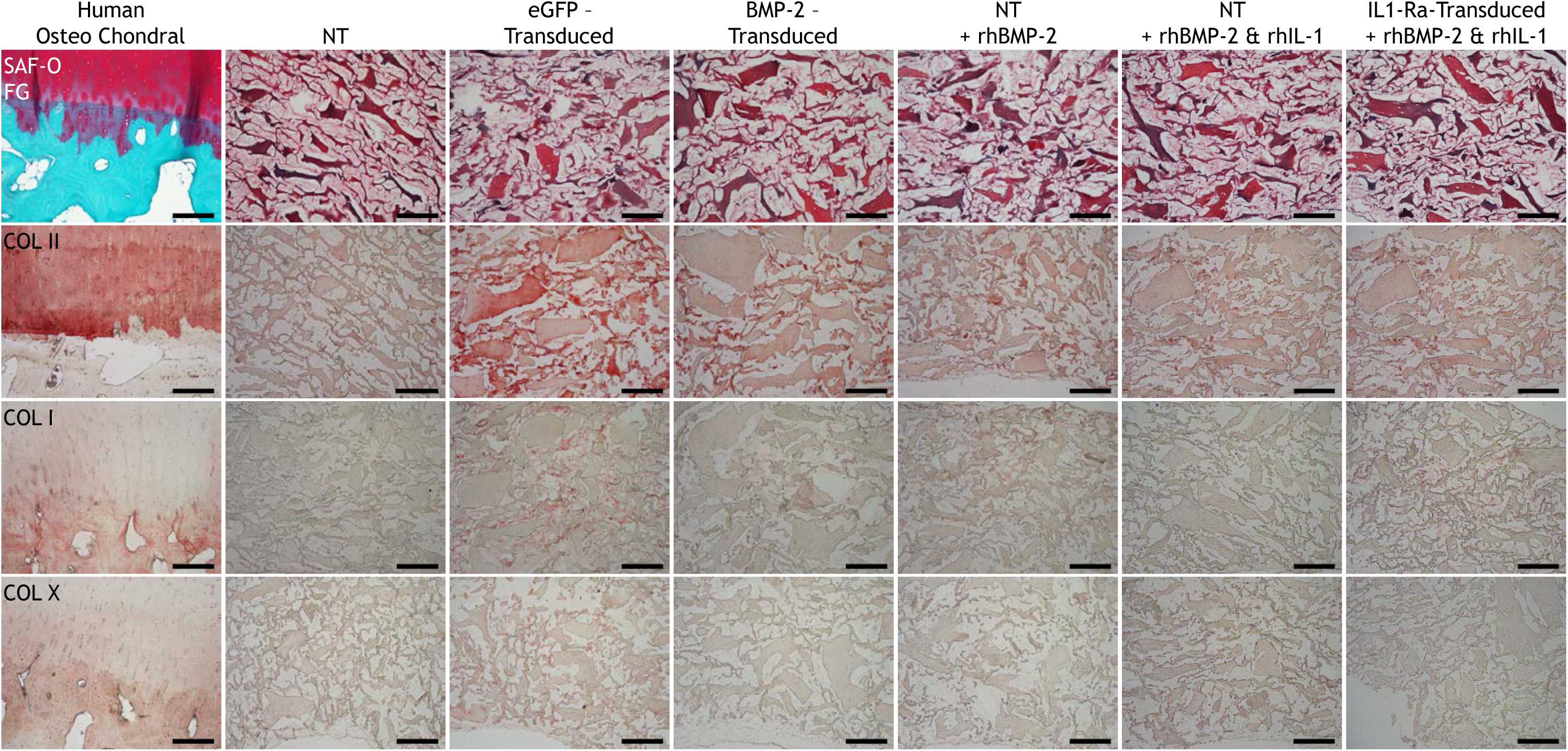
High magnification (10x) histology (Safranin-O and Fast Green staining); immunohistochemistry for type II collagen (COL II); type I collagen (COL I); type X collagen (COL X) on human osteochondral sections and non-transduced (NT), eGFP-transduced, or BMP-2-transduced constructs cultured in basal osteogenic media (without exogenous rhBMP-2), and NT constructs fed exogenous rhBMP-2 with and without rhIL-1, and IL-1Ra-transduced constructs fed exogenous rhBMP-2 and rhIL-1 after 28 days of culture in osteogenic media. All images are transverse sections, showing full-thickness of each construct. Aminoethyl carbazone (AEC) produces red color. Histology scale: 250μm.

**Supplemental Figure 8:**
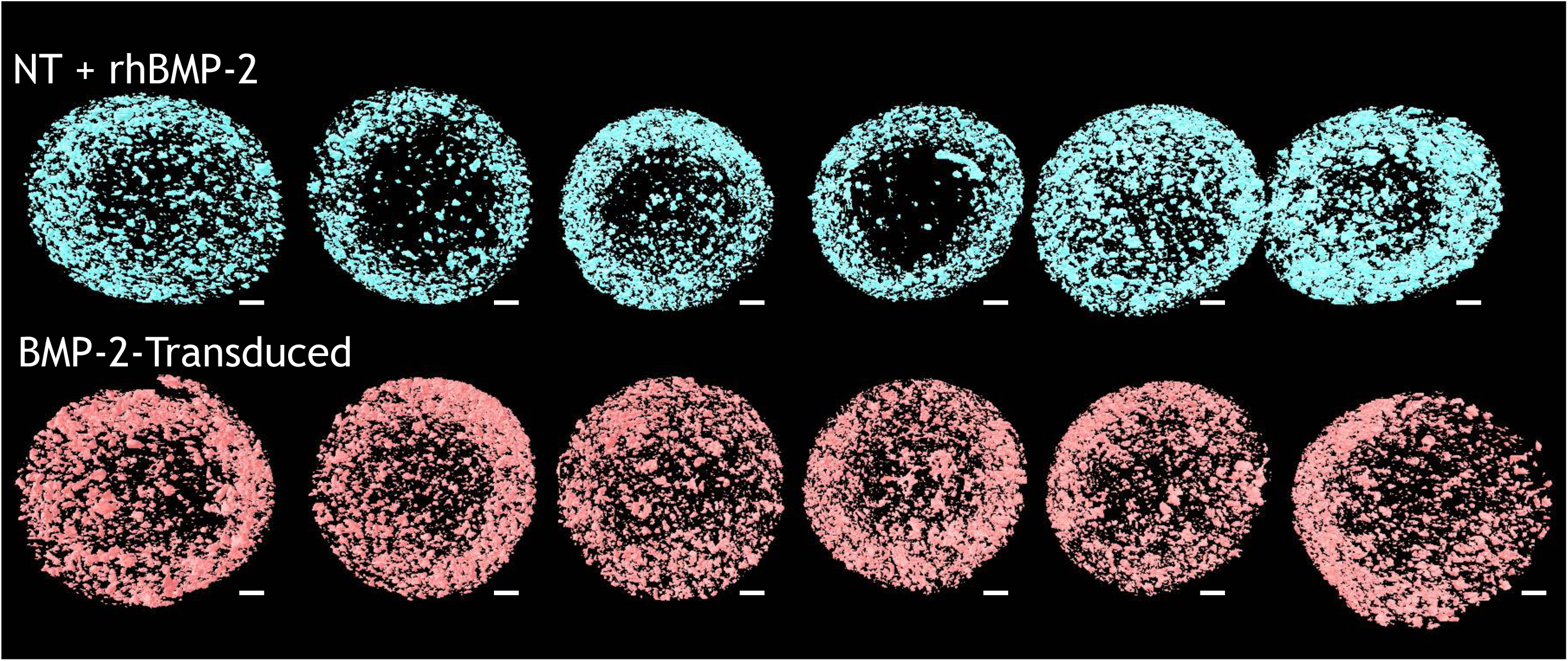
Micro-CT images of mineral distribution in (Top) non-transduced (NT) constructs fed exogenous rhBMP-2 or (Bottom) BMP-2-transduced constructs fed basal osteogenic media (without exogenous rhBMP-2). Image shows all 6 constructs per treatment group that were used to calculate the mineral distribution ratio presented in Figure 7E Scale: 1mm.

**Supplemental Video:**
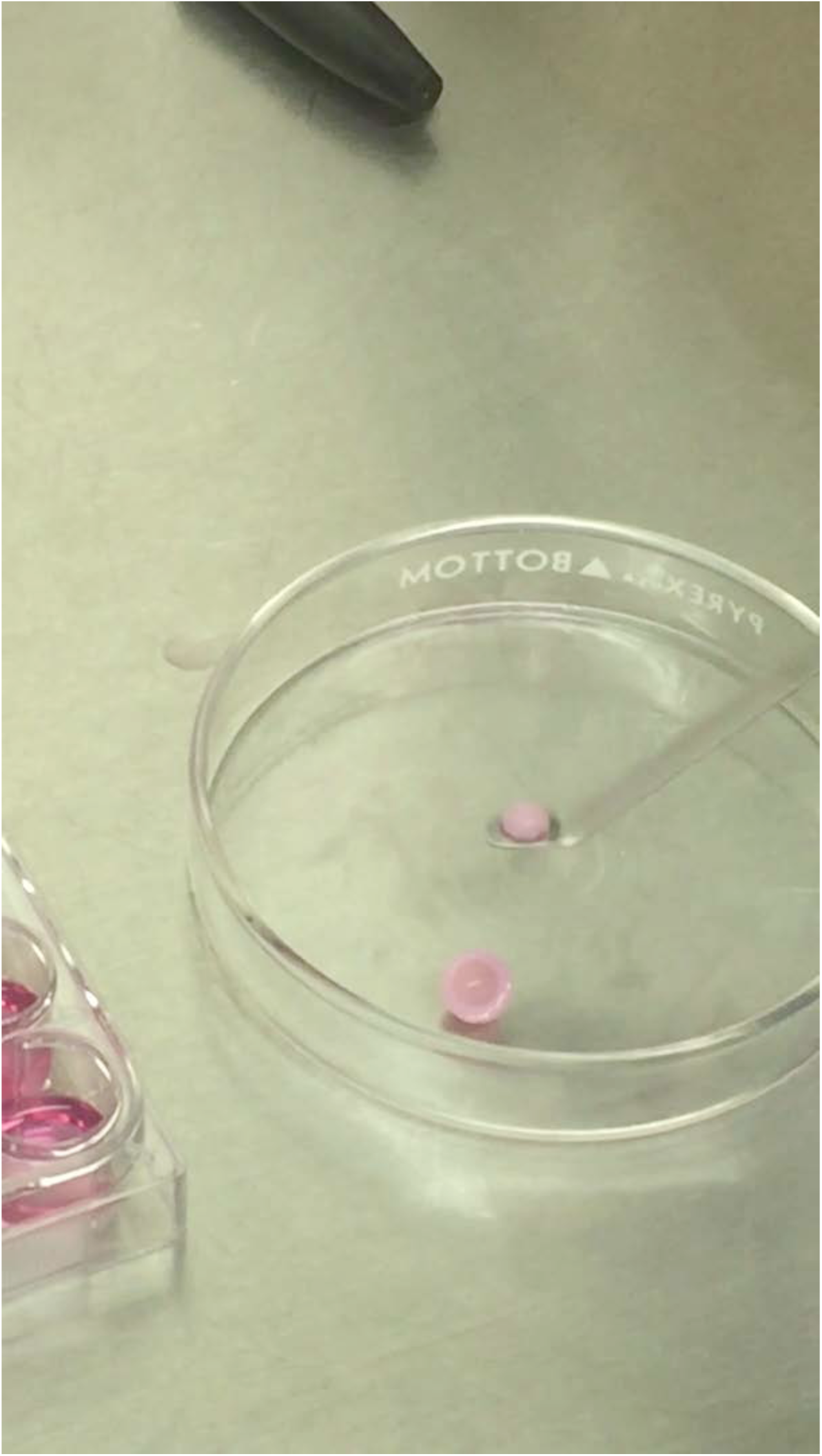
Assembly of osteochondral constructs by inserting inner hemisphere core into outer hemispherical shell after 22 days of respective osteogenic or chondrogenic differentiation.

